# B cell response to HCMV gH/UL116 elicits neutralizing and FcγR-activating antibodies

**DOI:** 10.64898/2026.10.11.758335

**Authors:** Artem Ashurov, Katja Hoffmann, Ajit R. Ramamohan, Krithika P. Karthigeyan, Anna Ashurov, Markus Rotzler, Ann-Kathrin Kohl, Gretel M. Torres, Christian Binuya, Caitlin Williams, Chelsea M. Crooks, Linda Schlachter, Danylo Shestopal, Leon Ullrich, Chiara Hornung, Ramin Lotfi, Philipp Kolb, Christoph Kreer, Manuel Koch, Sallie R. Permar, Hartmut Hengel, Jason S. McLellan, Florian Klein, Matthias Zehner

## Abstract

Humoral immune mechanisms that can contribute to limiting HCMV disease include neutralizing antibodies that block viral entry and Fc-mediated effector functions that promote elimination of infected cells. Here, we identify the gH/UL116 envelope complex as a naturally immunogenic target that elicits antibodies capable of mediating both of these immune mechanisms. Using single-cell sorting of antigen-specific memory B cells, we isolated gH/UL116-reactive monoclonal antibodies from HCMV-seropositive donors. Epitope mapping showed that gH binding was required for virus neutralization, whereas UL116 targeting was associated with potent and broad Fcγ receptor (FcγR) activation. Furthermore, cryo-electron microscopy provided high-resolution structures revealing how UL116-specific antibodies bind gH/UL116. In mice, gH/UL116 immunization elicited serum IgG responses whose binding titers strongly correlated with both neutralization and mouse FcγR activation. Together, these findings provide an epitope-level map linking antibody specificity to neutralization and FcγR activation across HCMV gH-associated complexes and position gH/UL116 as a promising HCMV vaccine candidate.

## INTRODUCTION

Human cytomegalovirus (HCMV) is an enveloped virus of the *Betaherpesvirinae* subfamily with a large double-stranded DNA genome (∼235 kb) that establishes lifelong persistence through latency following primary infection.^1,2^ Reactivation can occur during immunosuppression or inflammatory stress, and HCMV infection can cause severe or life-threatening disease in immunologically vulnerable populations, including fetuses and newborns, transplant recipients, and people living with advanced HIV infection.^1,2^ Congenital HCMV infection, a particularly severe manifestation as the leading infectious cause of birth defects and brain damage, affects approximately 0.5–1% of live births worldwide, with about 20% of infected infants developing clinical disease or permanent sequelae.^3,4^ Beyond overt disease, chronic HCMV infection has been associated with frailty and increased mortality in older adult cohorts.^5^ Despite this considerable burden, no HCMV vaccine is currently licensed, and available antivirals remain constrained by toxicity, the emergence of drug resistance, and limited efficacy.^6,7^ The development of safe and effective preventive and therapeutic strategies against HCMV therefore remains a high public health priority.^6–8^

The HCMV genome encodes approximately 750 translation products,^9^ including numerous membrane proteins.^10,11^ Entry into host cells is mediated by a coordinated interplay of viral envelope proteins, including the fusion glycoprotein B (gB) and receptor-binding gH/gL-associated complexes, the trimer (gH/gL/gO) and the pentamer (gH/gL/UL128-131A).^12–14^ These complexes engage distinct host receptors and together confer the remarkably broad cell tropism characteristic of HCMV.^15–17^ Beyond these canonical entry factors, other HCMV envelope proteins participate in virion assembly, egress, or modulate immune recognition, while many virion-associated proteins remain incompletely characterized.^18–21^ Among these is the noncanonical gH/UL116 complex (UL116 is hereafter termed gK), which forms independently of gL and is abundantly present on the virion surface.^22–24^ Within gH/gK, gK occupies a binding surface on gH that overlaps extensively with the gL-binding site, making gH/gK and gH/gL assemblies mutually exclusive.^22,24^ gK has also been shown to support the assembly, maturation, and virion incorporation of canonical gH/gL complexes.^23^ More recently, gH/gK was found to associate with UL141 to form the GATE complex, which enhances endothelial cell entry.^24^ These findings suggest that gH/gK may influence viral entry and host cell tropism and, together with the high sequence conservation of gK across clinical and laboratory-adapted HCMV strains,^22^ support evaluating gH/gK as a vaccine antigen and antibody target.^22–24^ However, naturally elicited humoral responses to gH/gK remain uncharacterized.

HCMV infection elicits broad humoral and cellular immune responses and can substantially reshape the composition and function of the immune system.^25–27^ Among humoral targets, B cell and antibody responses to gB and gH/gL-containing complexes have been extensively characterized, and antibodies against these proteins can potently neutralize HCMV *in vitro*.^28–31^ However, emerging evidence from cohort and vaccine studies suggests that antibody-associated protection is not fully explained by neutralization alone.^32–34^ Maternal Fc-dependent non-neutralizing antibody responses, including stronger antibody-dependent cellular cytotoxicity (ADCC) and antibody-dependent cellular phagocytosis (ADCP), were associated with a lower risk of congenital HCMV transmission.^33,34^ Functional studies have further demonstrated that human monoclonal antibodies can mediate ADCC against HCMV-infected cells.^35^ Collectively, this evidence supports a contributory role for FcγR-dependent antibody effector activity, alongside neutralization, in HCMV control. Nevertheless, systematic, epitope-resolved analyses of how antibodies targeting HCMV envelope complexes activate Fcγ receptors are lacking.

To address these gaps, we used single-cell sorting to isolate gH/gK-specific memory B cells from HCMV-seropositive human donors, demonstrating that gK is a target of the naturally elicited humoral immune response. We next analyzed the characteristics of the resulting monoclonal antibodies (mAbs) structurally and functionally, mapping epitopes by competition-based screening and resolving the binding sites of gH/gK-specific mAbs by cryo-electron microscopy. By assessing the antiviral functions of these mAbs, our study provides molecular insight into epitope-dependent patterns of neutralization and FcγR activation. Finally, we evaluated the immunogenicity of gH/gK and its ability to induce neutralizing and FcγR-activating antibodies in mice, informing its potential as a vaccine candidate.

## RESULTS

### HCMV gH/gK is a target of natural humoral immune response

To study the humoral immune response against HCMV gH/gK, we first expressed a soluble form of the complex (HCMV strain AD169) lacking the transmembrane domain of gH and assessed its purity and integrity by SDS-PAGE. As expected, gK migrated separately from gH due to the absence of an interprotein disulfide bridge, in contrast to the covalently linked gH/gL/gO trimeric complex (**Figure S1A**). To assess humoral reactivity to gH/gK, we isolated plasma IgG from four HCMV-seropositive blood donors (CS2– CS5) with broad and high neutralizing titers, selected from a larger cohort of 9,000 seropositive individuals^36^ (**Figure 1A**). Next, gH/gL-reactive antibodies were depleted from the isolated total plasma IgG of these donors, and residual binding to gH/gK was assessed to detect the gK-specific antibody response (**Figure 1B**). CS2 and CS4 showed high residual binding, indicating the presence of gK-specific antibodies in total plasma IgG, whereas CS5 showed lower remaining reactivity, and CS3 had no detectable gH/gK binding after gH/gL depletion (**Figure 1B**). To validate these findings, CS2 total plasma IgG was further tested using alternative depletion proteins. Depletion with gH/gL or gH/gL/gO left remaining IgG reactivity to gH/gK, confirming the presence of gK-specific antibodies in total plasma IgG (**Figure S1B**, blue curves). Interestingly, reactivity after gH/gL depletion was higher to gH/gK than to gH/gL/gO (**Figure S1B**), suggesting that total plasma IgG of CS2 contains quantitatively more antibodies targeting gK than those targeting gO.

**Figure 1.**
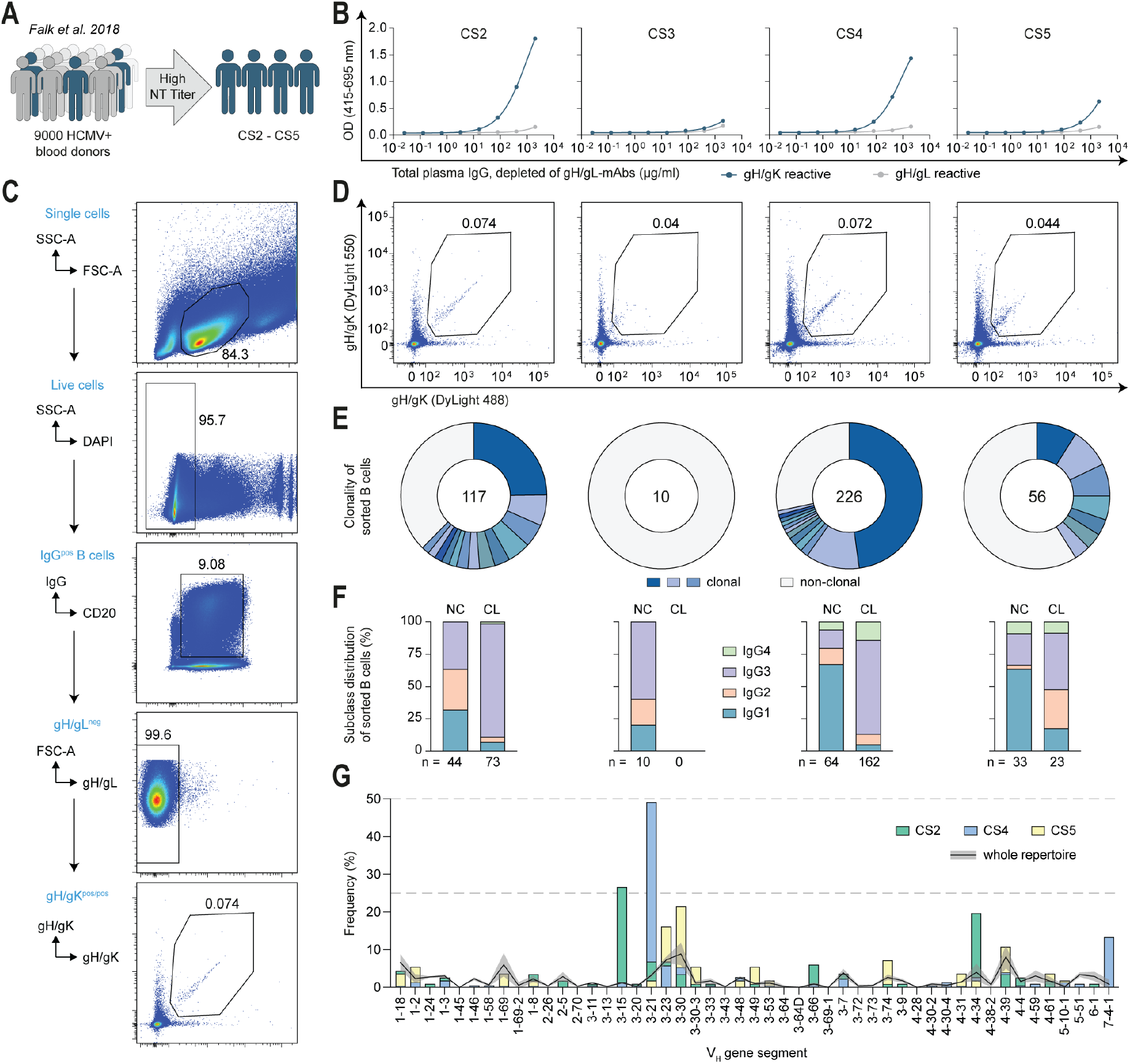
HCMV gH/gK is a target of natural humoral immune response. (**A**) Selection of top donors CS2–CS5 from a cohort of 9,000 HCMV-seropositive blood donors.^36^ Selection criteria were high neutralization potency in human foreskin fibroblasts (HFFs) and breadth across multiple HCMV strains.^36^ (**B**) gH/gK-specific residual binding (blue) measured by ELISA after depletion of total plasma IgG with gH/gL protein complex. Residual binding to gH/gL after gH/gL depletion is shown as a depletion control (gray). Representative data of at least two separate experiments is shown. OD, optical density. (**C**) Fluorescence-activated cell sorting (FACS) gating strategy for single-cell sorting of DAPI^−^, IgG^+^, gH/gL^−^ and gH/gK^+/+^ memory B cells, shown for a representative donor (CS2). (**D**) Sorting gates for IgG^+^ memory B cells across all donors, with the percentage of gH/gK^+/+^ cells indicated in each plot. (**E**) Clonality of isolated and sequenced B cells (defined as shared V_H_ and J_H_ gene usage and >75% CDRH3 amino acid similarity) shown as donut charts across all donors. Blue/green slices denote clonal groups; white slices denote non-clonal cells within the sorted population. Numbers in the centers indicate the count of successfully recovered B cell receptor sequences. (**F**) IgG-subclass distribution of heavy chain constant regions of sequenced B cells across all donors, comparing clonally related (CL) versus non-clonal (NC) B cells. (**G**) V_H_ gene distribution among sorted gH/gK-reactive B cells from donors CS2, CS4, and CS5 (CS3 was excluded due to low cell count). Colored bars show the frequency of specific V_H_ genes in the sorted repertoire for each donor. The black line indicates the mean V_H_ frequency in whole IgG-positive B cell repertoires from the same donors (determined by NGS), with gray shading denoting the standard deviation (SD) for each V_H_ gene.

To analyze the memory B cell response against gH/gK in these four donors on a single cell level, we sorted IgG^pos^ memory B cells that were gH/gL-negative and gH/gK double-positive, thereby enriching for gK-reactive cells and reducing the dominant gH-specific population (**Figure 1C and S1C**). This sorting strategy revealed a distinct gH/gK^pos/pos^ B cell population in CS2, CS4, and partly in CS5, whereas in CS3 only sporadic double-positive events were detected (**Figure 1D**). Notably, this aligns with the lower activity of gK-specific plasma IgG in CS5 and its absence in CS3 (**Figure 1B**), indicating that the humoral immune response to gH/gK is similarly represented across plasma IgG and memory B cell compartments.

We then sequenced the variable regions of the B cell receptors of sorted B cells and assessed their clonal relationships, defining clonally related B cells by shared heavy chain variable (V_H_) and joining (J_H_) gene usage and >75% amino acid similarity of the third complementarity determining region of heavy chains (CDRH3) (**Figure 1E**). Except for CS3, all other donors showed polyclonal responses with clonally-related sequences comprising 62% (CS2), 72% (CS4) and 41% (CS5) of sorted cells (**Figure 1E**). Several clones were expanded, with the largest in CS4 comprising 108 cells (48% of all sorted cells of CS4; **Figure 1E**). Of note, IgG subclass analysis of heavy chain constant regions revealed a bias toward IgG3 among clonally related cells (predominantly in CS2 and CS4; **Figure 1F**), whereas non-clonal cells showed no such IgG3 enrichment (**Figure 1F**). Next, we analyzed V_H_ gene usage of the isolated B cells across individuals and compared it to NGS-derived whole B cell repertoire of all donors (**Figure 1G**). This revealed donor-specific V_H_-gene preferences in gH/gK recognition, with IGHV3-15 and IGHV4-34 elevated in CS2 and IGHV3-21 overrepresented in CS4 (**Figure 1G**). V_H_ gene germline identity in sorted cells mirrored whole repertoires in CS2 and CS5, while it was skewed towards lower identities in CS4 (median 90.2% vs. 93.9%; **Figure S1D**), largely driven by an expansion of a dominant clone. Finally, the amino acid lengths of CDRH3, which are often key determinants of antigen binding, were on average slightly longer in CS2 and CS4 relative to their whole repertoires (**Figure S1E**), suggesting antigen-driven selection for longer paratopes in gH/gK-reactive clones in these donors.

Together, these data demonstrate that gK is a natural target of human humoral immunity, with gH/gK-reactivity detected both in plasma IgG and in the memory B cell compartments across multiple donors.

### Binding assays identify gH/gK-specific and gH/gL-cross-reactive mAbs

To assess their binding activity, we cloned B cell receptors from selected gH/gK^pos/pos^ B cells into secreting plasmids and produced 96 mAbs as human IgG1 antibodies in HEK293-6E cells. Selection prioritized representation of distinct B cell clones, inclusion of unique sequence variants within clones, and a focus on highly mutated B cells as indicated by lower V_H_ germline identity. Of the mAbs produced, 36 bound gH/gK by ELISA at a maximal tested concentration of 30 µg/ml (**Figure 2A**). To identify gK-specific antibodies, we next tested the reactivity of the gH/gK-binding mAbs against the gH/gL complex. Most gH/gK binders did not cross-react with gH/gL, whereas 12 showed detectable binding (**Figure 2A**, blue segments), indicating that the negative gH/gL gate during sorting was not fully exclusive. In CS3, the single binding mAb was cross-reactive to gH/gL; in CS5, three binding mAbs were detected, one of which was gH/gK-specific. The remaining binders were derived from CS2 and CS4 donors (**Figure 2A**), in concordance with their stronger total plasma IgG reactivity to gH/gK and higher frequencies of gH/gK-reactive B cells (**Figure 1**).

**Figure 2.**
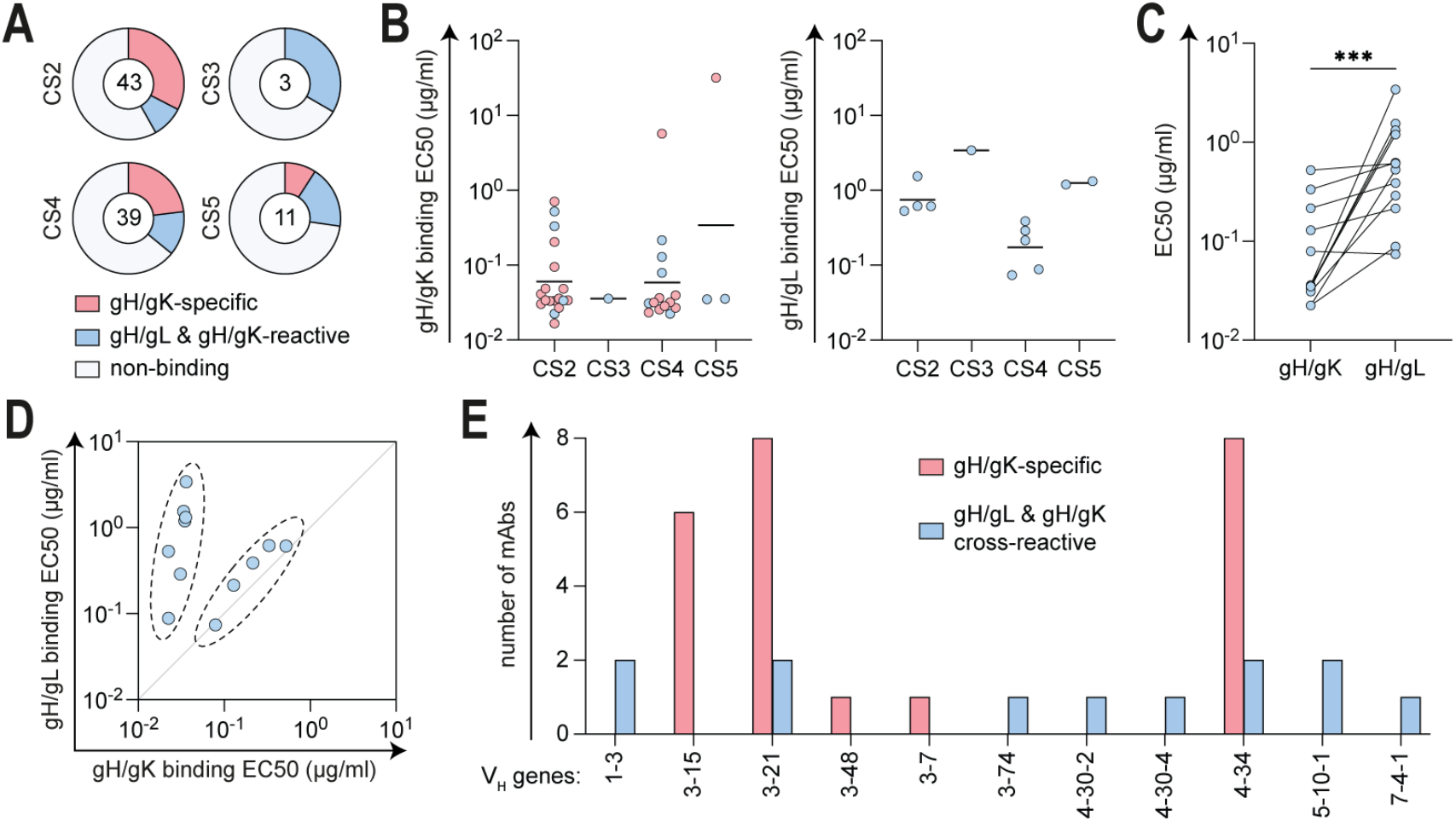
Binding assays identify gH/gK-specific and gH/gL-cross-reactive mAbs. (**A**) Overview of mAbs isolated from each donor. The number in the center of each donut plot indicates the total number of mAbs expressed and purified from HEK293 cells. Pink segments indicate mAbs that reacted only with gH/gK by ELISA, blue segments indicate mAbs that reacted with both gH/gK and gH/gL protein complexes, and white segments indicate mAbs that did not bind either gH/gK or gH/gL at the highest tested concentration of 30 µg/ml. (**B**) ELISA-derived EC_50_ values for binding to gH/gK (left) or gH/gL (right) complexes, separated by donor. Each dot represents one mAb and shows the geometric mean of at least two separate experiments. Black lines indicate geometric means of all mAbs isolated from each donor. (**C**) Comparison of EC_50_ values of cross-reactive mAbs for binding to gH/gK and gH/gL protein complexes. Values for the same mAb are connected by lines. Significance was assessed using a two-tailed Wilcoxon matched-pairs signed-rank test. Data are derived from Figure 2B and include only cross-reactive mAbs. (**D**) 2D plot of EC_50_ values of cross-reactive mAbs, with gH/gK binding EC_50_ values shown on the X-axis and gH/gL binding EC_50_ values shown on the Y-axis. Dashed outlines indicate two visually apparent groups of cross-reactive mAbs based on EC_50_ values: mAbs with comparable binding to both antigens (right group), and mAbs with preferential binding to gH/gK (left group). Data are derived from Figure 2B and include only cross-reactive mAbs. (**E**) V_H_ gene distribution among gH/gK-specific mAbs (pink) and gH/gL-cross-reactive mAbs (blue). Bars indicate the number of mAbs using respective V_H_ gene.

ELISA-determined EC_50_ values indicated strong binding for most gH/gK-reactive mAbs, ranging from 0.02 to 0.71 µg/ml, with two higher EC_50_ outliers (**Figure 2B**, left). Binding to gH/gL was generally weaker, with the lowest EC_50_ of 0.07 µg/ml (**Figure 2B**, right). A direct comparison showed lower average EC_50_ values of cross-reactive mAbs to gH/gK than to gH/gL (**Figure 2C**). Notably, when EC_50_ values for both antigens were plotted against each other, the mAbs separated into two groups: one with comparable binding to both antigens (**Figure 2D**, right dashed outline) and another with preferential binding to gH/gK over gH/gL (**Figure 2D**, left dashed outline). This binding pattern suggests recognition of distinct epitopes on the gH/gK surface by cross-reactive mAbs.

Finally, we analyzed V_H_ gene usage among binders and found that while cross-reactive mAbs were distributed across different V_H_ genes, gH/gK-specific mAbs were enriched for IGHV3-15, IGHV3-21, and IGHV4-34 genes (**Figure 2E**), consistent with the sorting results.

We conclude that isolated gH/gK-reactive memory B cells encode mAbs with strong gH/gK binding, including both gH/gK-specific mAbs and mAbs cross-reactive with gH/gL.

### Neutralization by gH/gK-targeting mAbs requires engagement of gH

To classify the epitopes of gH/gK-reactive mAbs, we selected a clonally representative subset of 22 mAbs, including at least one representative from each B cell clonal group among the binders, and aimed to group this panel by competition ELISA. To this end, mAbs were biotinylated and used as competitors against the same unlabeled mAbs, allowing assignment into four competition groups (**Figure 3A**). Group 1 contained both gH/gK-specific and gH/gL cross-reactive mAbs, indicating the recognition of an overlapping epitope near the gH/gK interface. Groups 2 and 3 comprised gH/gK-specific mAbs, while group 4 split into subgroup 4a with three gH/gL cross-reactive mAbs and subgroup 4b with one cross-reactive mAb (**Figure 3A**). Notably, competition patterns for groups 3 and 4a showed a substantial overlap despite differing specificities (**Figure 3A**), suggesting steric interference or epitope occlusion due to conformational effects.

**Figure 3.**
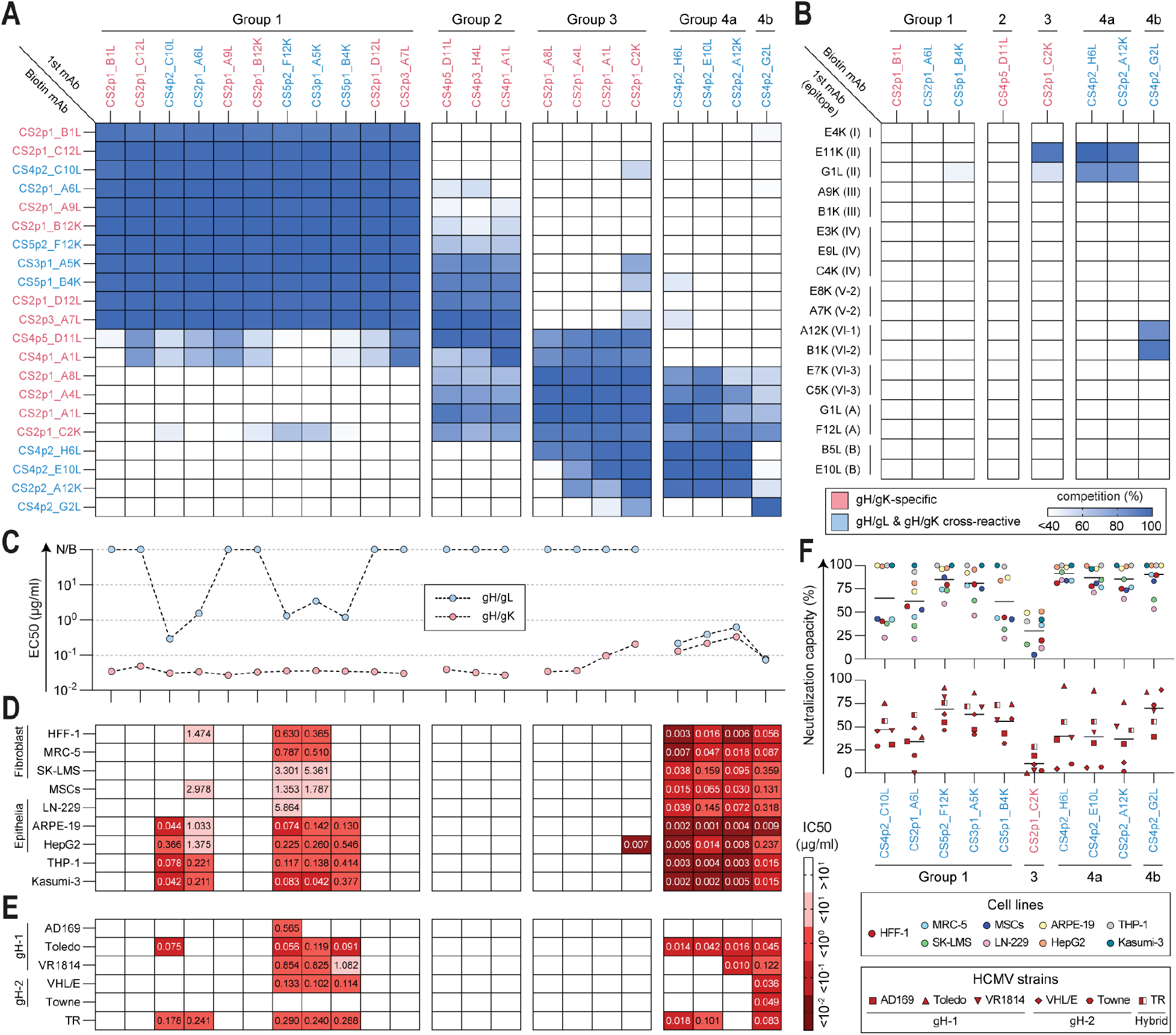
Neutralization by gH/gK-targeting mAbs requires engagement of gH. (**A**) Competition ELISA for binding to the gH/gK complex using biotinylated gH/gK-targeting mAbs (rows) and unlabeled mAbs (columns). Competition is shown in blue. gH/gK-specific mAbs are labeled in pink, cross-reactive mAbs are labeled in blue. mAbs are clustered according to their competition patterns. (**B**) Competition ELISA for binding to the gH/gK complex using representative mAbs from the groups identified in Figure 3A tested against a reference panel of 18 gH/gL-targeting mAbs.^31,37^ Reference mAbs are grouped according to previously described epitope groups,^31,37^ with group assignments indicated in parentheses next to each reference mAb. Competition is shown in blue. gH/gK-specific mAbs are labeled in pink, cross-reactive mAbs are labeled in blue. (**C**) ELISA-derived EC_50_ values for binding to gH/gK (pink) or gH/gL (blue) complexes. mAbs are shown in the same order as in Figure 3A, with values for mAbs belonging to the same competition group connected by dashed lines. Each dot shows the geometric mean of at least two separate experiments. Data are derived from Figure 2B. N/B, non-binding. (**D**) Neutralizing activity of gH/gK-targeting mAbs against HCMV_TB40/E_ infection in different host cell lines, shown as IC_50_ values (µg/ml) in a sequential red color scale. Host cell lines included fibroblast-like cells (HFF-1, MRC-5, SK-LMS-1), epithelial-like cells (LN-229, ARPE-19, HepG2), mesenchymal stem cells (MSCs), monocytes (THP-1), and myeloid cells (Kasumi-3). Each IC_50_ value was calculated from two separate experiments. (**E**) Neutralizing activity of gH/gK-targeting mAbs against infection with different HCMV strains in HFF-1 fibroblasts, shown as IC_50_ values (µg/ml) in a sequential red color scale. HCMV strains were selected based on gH genotype: gH-1 (AD169, Toledo, VR1814), gH-2 (VHL/E, Towne), and hybrid gH (TR). Each IC_50_ value was calculated from two separate experiments. (**F**) Neutralizing capacity of gH/gK-targeting mAbs against HCMV_TB40/E_ infection in different host cell lines (top) and against infection with different HCMV strains in HFF-1 fibroblasts (bottom). Neutralizing capacity was calculated from mAb dose-response curves and represents the difference between the infection rate in no-antibody conditions and the residual infection plateau at saturating mAb concentrations (also see Methods). Each value was calculated from two separate experiments.

To further define these epitopes, we tested representative mAbs from each group by competition ELISA against a panel of 18 reference mAbs that target distinct, previously defined and structurally characterized sites on gH/gL.^31,37^ Multiple reference mAbs from this panel interact with gH and were able to bind the gH/gK complex (**Figure S2A**). Whereas group 3 and group 4 mAbs competed with some of the reference mAbs, group 1 and group 2 did not overlap with any mAbs from the gH/gL reference panel (**Figure 3B**). These findings suggest that group 1 and 2 mAbs recognize epitopes located either on gK or within gH regions not covered by reference mAbs.

Consistent with the EC_50_-based grouping observed by ELISA (**Figure 2D**), group 4 mAbs bind gH/gL and gH/gK with similar EC_50_ values, whereas group 1 mAbs show lower EC_50_ to gH/gK (**Figure 3C**). These relationships suggest that group 4 mAbs recognize accessible gH epitopes present in both complexes, while group 1 cross-reactive mAbs target epitopes preferentially presented in the gH/gK complex.

Next, we evaluated neutralizing activity of grouped mAbs across HCMV-susceptible cell lines, including various epithelial cell types, fibroblasts, mesenchymal stem cells, monocytes, and myeloid cells. Importantly, neutralization was mediated almost exclusively by cross-reactive mAbs (**Figure 3D**). Group 4 mAbs potently neutralized HCMV infection in all tested cell types, with IC_50_ values ranging from 0.001 to 0.359 µg/ml (**Figure 3D**). Cross-reactive group 1 mAbs showed more moderate activity, with IC_50_ values ranging from 0.042 to 5.864 µg/ml, and some failed to neutralize HCMV infection in individual cell types (**Figure 3D**). Notably, several group 4 mAbs reached potencies comparable to reference mAb CS4tt1p1_E3K (referred to as E3K), a gH/gL-specific broadly neutralizing mAb characterized previously^31^ (**Figure S2B**). Interestingly, the gH/gK-specific mAb CS2p1_C2K yielded a detectable IC_50_ in HepG2 hepatocytes (0.007 µg/ml), whereas neutralization in other cell types did not reach the 50% threshold required for IC_50_ determination (**Figure 3D**). Additionally, we tested all mAbs against infection of HFF-1 fibroblasts by different HCMV strains representing distinct gH genotypes (gH-1: AD169, Toledo, VR1814; gH-2: VHL/E, Towne; hybrid: TR). The same pattern persisted, with neutralization restricted to cross-reactive mAbs (**Figure 3E**). Unlike the multi-cell-line panel, group 4 mAbs did not neutralize all strains to the same extent (**Figure 3E**). This limited neutralization reflects a reduced neutralizing capacity of mAbs and strain-dependent differences: while all group 4 mAbs nearly completely neutralized TB40/E strain (gH-1 genotype) across cell lines (**Figure 3F**, top), their activity against most other strains resulted in higher residual infection levels (**Figure 3F**, bottom). Group 1 cross-reactive mAbs likewise did not achieve complete neutralization, approaching close to 100% neutralization capacity only in monocytes, myeloid cells, and some epithelial cell lines (**Figure 3F**).

Together, these data indicate that neutralization by gH/gK-targeting mAbs requires engagement of gH, with potency largely determined by epitope location.

### FcγR activation by gH/gK-targeting mAbs is epitope- and subclass-dependent

We next tested whether gH/gK-reactive mAbs recognize their targets displayed on the surface of infected cells. Five days post-infection of fibroblasts with HCMV, staining with gH/gK-specific CS2p1_B1L or cross-reactive CS4p2_G2L produced comparable surface signals, similar to gH-targeting reference mAbs G1L and E10L^37^ (**Figure 4A**, left panels). In contrast, mAbs specific to gH/gL, gO, or UL128-131A yielded much weaker signals (**Figure 4A**, right panels). These data show that gH/gK-reactive mAbs recognize native gH/gK complex on the surface of infected cells and suggest that, at five days post-infection, gH/gK is more abundant on infected-cell surfaces than other gH-associated complexes.

**Figure 4.**
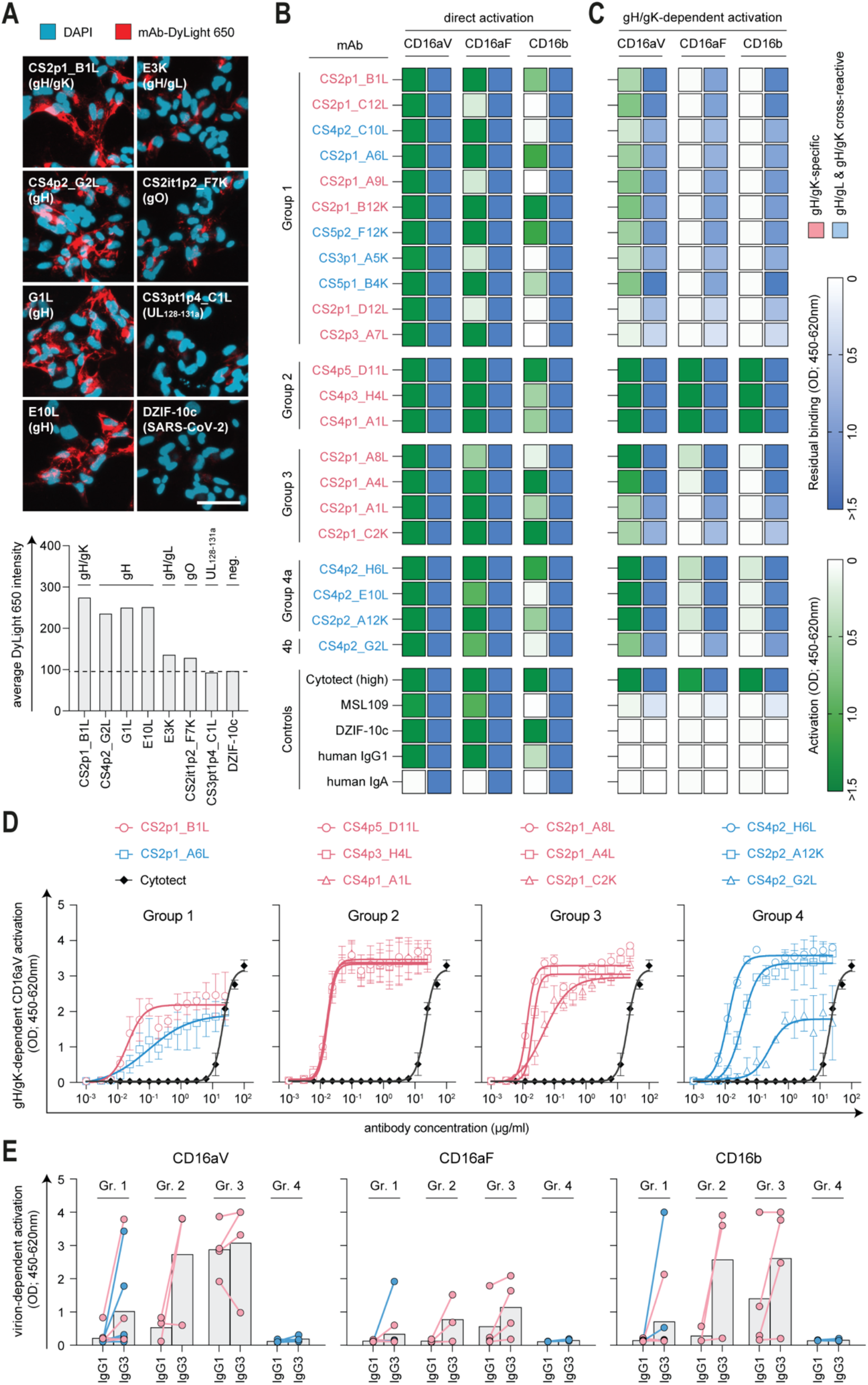
FcγR activation by gH/gK-targeting mAbs is epitope- and subclass-dependent. (**A**) Staining of HCMV proteins expressed on the cell surface of HFF-1 fibroblasts 5 days post infection with HCMV_TB40/E_. For surface staining, infected cells were first incubated with the purified human non-HCMV specific antibody DZIF-10c^39,40^ (SARS-CoV-2-specific mAb) to block HCMV-encoded Fc receptors^41^ and subsequently stained with DyLight 650-labeled mAbs. Staining was performed before fixation to limit intracellular mAb access (also see Methods). Individual panels show staining with the indicated mAbs, with mAb specificity indicated in parentheses. All panels are shown with the same brightness and contrast settings and are representative of each mAb staining. Scale bar = 50 μm. Quantification below the microscopy panels shows the average cellular pixel intensity of the DyLight 650-positive signal for each mAb. The dashed line indicates the average pixel intensity of the DZIF-10c negative control mAb^39,40^ for comparison. (**B–C**) Human CD16 activation by gH/gK-reactive mAbs measured using the BW5147 reporter cell system under antigen-independent mAb coating conditions for direct FcγR crosslinking and activation (Figure 4B), or under antigen-dependent conditions with mAbs bound to coated gH/gK complex (Figure 4C). For each mAb, condition, and CD16 variant (CD16aV, CD16aF, and CD16b), paired heatmap squares show CD16 activation in green and residual antibody binding in blue. Activation was measured by ELISA-based IL-2 detection in supernatants of the respective reporter cells; residual binding of antibodies remaining after each activation readout was measured by ELISA within the same assay as a loading control. gH/gK-specific mAbs are indicated by pink text labels, and cross-reactive mAbs by blue text labels. mAbs are arranged by epitope group, with control antibodies shown below. mAbs were used at 1 μg/ml, Cytotect was used at 10 μg/ml. Each antibody/condition was tested in at least two separate experiments; representative data from one experiment are shown. (**D**) Human CD16aV activation measured using the BW5147 reporter cell system under gH/gK complex coating conditions. Antibodies were serially diluted to assess dose–response characteristics of CD16aV activation, measured by ELISA-based IL-2 detection. mAbs are arranged by epitope group, with up to three representative mAbs tested from each epitope group. The symbol key for mAbs is indicated above, and Cytotect is shown in all plots as black curves. Mean values and SDs from two separate experiments are shown. (**E**) Human CD16 activation by gH/gK-reactive mAbs, measured using BW5147 reporter cells expressing CD16aV, CD16aF, or CD16b under virion-dependent conditions with immobilized HCMV_TB40/E_ virions. Activation was measured by ELISA-based IL-2 detection in supernatants of the respective reporter cells. Dots show activation by individual mAbs, with lines connecting the IgG1 and IgG3 measurements for the same mAb. Pink and blue dots indicate gH/gK-specific and gH/gL-cross-reactive mAbs, respectively. mAbs are arranged by epitope group, and gray bars indicate mean activation for each group and IgG subclass. mAbs were used at 50 μg/ml. Each antibody was tested in at least three separate experiments; representative data from one experiment are shown.

To evaluate Fc gamma receptor (FcγR) crosslinking and activation capability by gH/gK-reactive mAbs, we used the BW5147 reporter cell system^38^ to perform CD16 (FcγRIII) activation assays using different coating conditions (**Figure 4B–C and S3A**). All mAbs were expressed with the same human IgG1 Fc region, allowing differences in FcγR activation to be assessed independently of Fc sequence variation. In the antigen-independent direct activation assay, all mAbs activated CD16aV (the high-affinity CD16a variant), confirming intact Fc presentation upon mAb immobilization on the plate (**Figure 4B**). Interestingly, some mAbs did not activate the lower-affinity CD16aF (**Figure 4B**, green boxes), despite being present on the plate after the readout (**Figure 4B**, blue boxes). CD16b was likewise not uniformly activated, consistent with its low affinity for IgG (**Figure 4B**). A lack of CD16aF or CD16b activation by certain mAbs under antigen-independent conditions may reflect differences in receptor affinity and activation thresholds as well as intrinsic variation in the FcγR-activating potential among mAbs.

gH/gK antigen-dependent FcγR activation assays revealed epitope-specific differences in CD16 activation: groups 1 and 3 mAbs activated only CD16aV and did so at lower levels than in the antigen-independent format (**Figure 4C**, green boxes), consistent with reduced mAb presentation upon binding to gH/gK (**Figure 4C**, blue boxes). Group 4 mAbs activated CD16aV but showed markedly lower CD16aF and CD16b activation on gH/gK complex despite robust antigen binding (**Figure 4C**), indicating that antigen binding alone is not sufficient for efficient receptor activation. In contrast, gH/gK-specific group 2 mAbs broadly activated all CD16 variants to a high level, comparable to HCMV-reactive polyclonal immunoglobulin (Cytotect) that was tested at tenfold higher concentration and used as a positive control (**Figure 4C**). Notably, gH/gL/gO antigen-dependent assays further confirmed that the broad CD16 activation pattern of group 2 mAbs is gH/gK-restricted, with no detectable binding or FcγR activation mediated by gH/gL/gO complex (**Figure S3A**).

In addition, we evaluated the CD16 activation properties of mAbs from the reference panel^31,37^. In contrast to gH/gK-reactive mAbs, only a subset of gH/gL-targeting reference mAbs activated CD16aV, with weaker and less sustained activation of CD16aF and CD16b (**Figure S3B**). Other gH/gL-targeting mAbs showed little to no CD16 activation, despite antigen-independent direct activation potential and detectable residual binding to gH/gL/gO for most mAbs (**Figure S3B**).

We further assessed the potency of FcγR activation by gH/gK-reactive mAbs by testing representative mAbs from each epitope group in a dose–response CD16aV activation assay in gH/gK antigen-dependent conditions (**Figure 4D**). All tested mAbs showed markedly greater CD16aV activation potency than polyclonal Cytotect, with EC_50_ values up to three orders of magnitude lower (**Figure 4D**). Notably, while most mAbs reached maximal effects comparable to Cytotect, group 1 and group 4b mAbs plateaued at lower OD values (**Figure 4D**), further supporting epitope-dependent differences in FcγR activation by gH/gK-reactive mAbs.

To extend our analysis to virion-associated antigens, we next tested CD16 activation by gH/gK-reactive mAbs using immobilized HCMV_TB40/E_ virions (**Figure 4E**). Initial experiments, performed with the human IgG1 mAbs used in the preceding assays, revealed little or no CD16 activation for most antibodies, with responses largely restricted to group 3, particularly for CD16aV and CD16b (**Figure 4E**). Given that most isolated mAbs originated from IgG3-expressing memory B cells (**Figure 1F**), we next tested the same antibodies with a human IgG3 backbone. This substantially enhanced activation by several mAbs, with increases observed for all CD16 variants (**Figure 4E**). These gains were particularly pronounced for gH/gK-specific mAbs of group 2 and selected group 1 mAbs. In contrast to their activity with recombinant proteins, cross-reactive group 4 mAbs induced minimal CD16 activation in the virion-based assay in either the IgG1 or IgG3 format (**Figure 4E**).

Finally, we assessed virion-dependent activation of human CD32aH, CD32aR, CD32b, and CD64 reporter cells, comparing gH/gK-reactive mAbs in the IgG1 and IgG3 formats (**Figure S3C**). CD32 reporter activation broadly resembled the pattern observed for CD16, with gH/gK-specific group 2 and 3 mAbs eliciting higher mean responses than the other groups. Group 2 responses were largely restricted to the IgG3 format, whereas group 3 mAbs induced activation in both subclass formats (**Figure S3C**). Selected group 1 mAbs also showed strong CD32 reporter activation in the IgG3 format, while group 4 mAbs remained weakly active. CD64 reporter responses were observed across all epitope groups in both formats, including modest responses from group 4 mAbs. Mean responses for group 1 approached those of group 2, whereas group 3 showed the highest mean activation. Interestingly, differences between the IgG1 and IgG3 formats were less pronounced for CD64 (**Figure S3C**).

We conclude that gH/gK-targeting mAbs show epitope- and subclass-dependent differences in FcγR activation, with gH/gK-specific epitopes generally supporting broader and stronger responses than epitopes recognized by gH/gL-cross-reactive mAbs in the virion-based assays.

### Cryo-EM structures reveal gH/gK-specific antibody epitopes

To define the molecular epitopes on the gH/gK complex associated with broad FcγR activation, we next sought to determine cryo-EM structures of gH/gK in complex with selected antibodies. We prioritized group 2 because its mAbs showed broad CD16 activation with recombinant gH/gK (**Figure 4C–D**) and, in the IgG3 format, strong FcγR activation in virion-based assays (**Figure 4E and Figure S3C**). We selected CS4p5_D11L (hereafter, D11L) as a representative mAb from this group. Based on the competition patterns (**Figure 3A**), we also included the group 1 mAb CS2p1_B1L (hereafter, B1L) and the group 4b mAb CS4p2_G2L (hereafter, G2L) for combined cryo-EM analysis, as these mAbs did not compete with D11L or with each other. We determined a 3.3 Å resolution cryo-EM structure of gH/gK bound to the antigen-binding fragments (Fabs) of all three mAbs (**Figure S4 and Figure S5**). Because D11L Fab displayed a flexible binding orientation relative to gH/gK, 3D classification followed by local refinement was required to resolve its interaction with the complex, yielding a 3.5 Å resolution local reconstruction (**Figure S4 and Figure S5**). The Fabs recognized distinct, non-overlapping epitopes: G2L bound a membrane-proximal epitope on gH, B1L bound a quaternary epitope bridging gH and gK, and D11L bound an epitope located predominantly on gK (**Figure 5A**).

**Figure 5.**
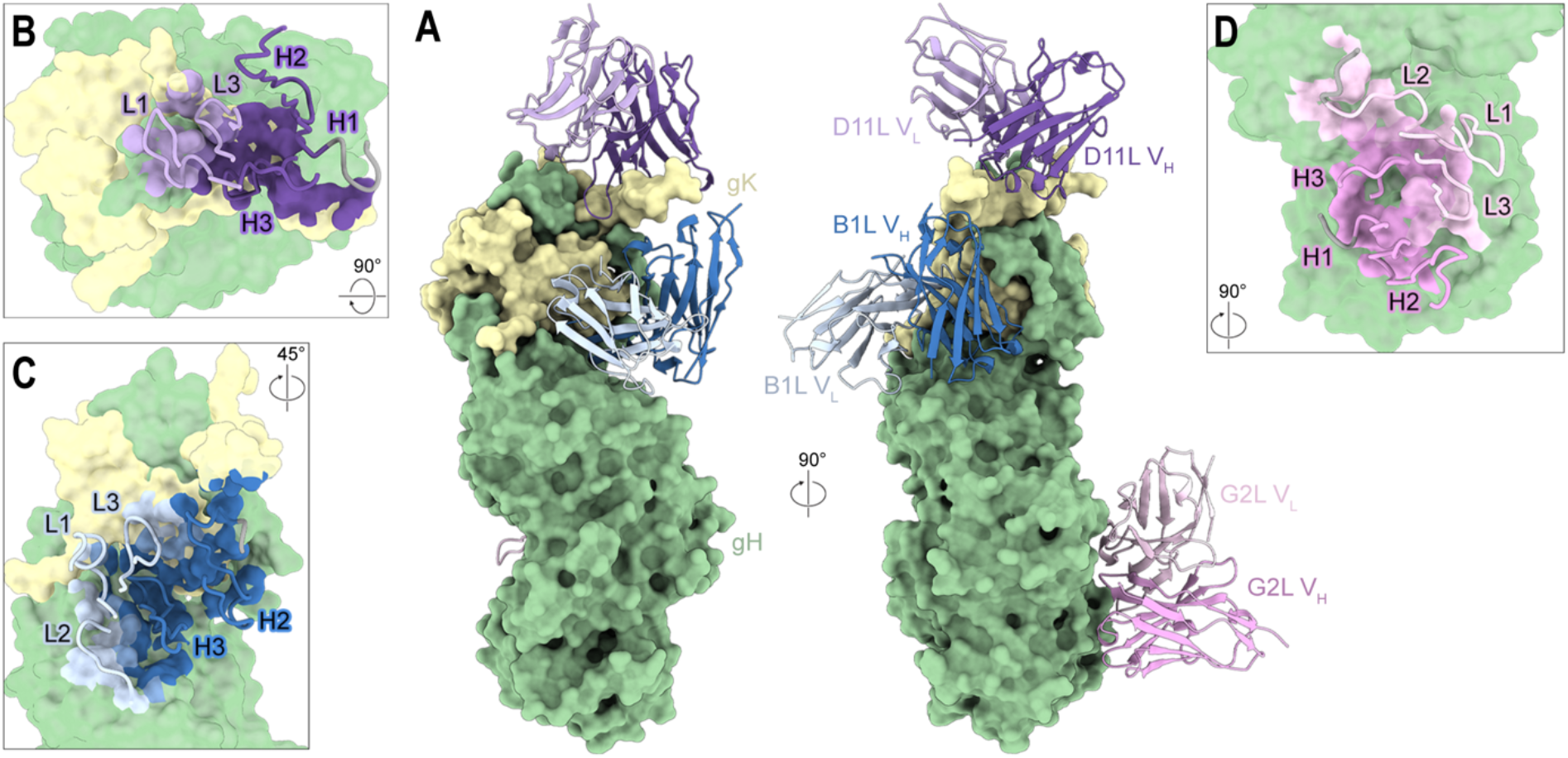
D11L, B1L, and G2L bind three non-overlapping epitopes across gH/gK. (**A**) Two side views of gH/gK bound by D11L, B1L, and G2L Fabs, rotated by 90 degrees. Antibody chains shown as cartoons, with opaque surfaces for gH (green) and gK (yellow). (**B–D**) Epitopes of D11L (Figure 5B), B1L (Figure 5C), and G2L (Figure 5D), respectively. Surface views of gH/gK with epitopes colored by-atom. CDR loops are shown as noodles, with additional framework residues shown in gray.

The epitope of G2L lies entirely on gH (**Figure 5A**, right), consistent with its classification by competition ELISA (**Figure 3A**), and buries approximately 1040 Å^2^ of gH surface area. The epitope is markedly discontinuous, assembled from four widely separated segments of the gH sequence — residues ∼433– 434, ∼479–483, ∼568–588, and ∼620–626, with an additional contact near 701 — that are brought together in space at the membrane-proximal region of gH (**Figure 5A**). G2L engages the antigen through all three CDRs of both chains (**Figure 5D**). Interestingly, superposition with the recently described GATE complex^24^ revealed that G2L binding would sterically clash with UL141 (**Figure S6A**). As the GATE complex has been shown to enhance HCMV entry into endothelial cells,^24^ interference with its assembly or function may contribute to the neutralizing activity of G2L.

B1L forms the largest interface of the three Fabs, burying approximately 1650 Å^2^ of surface area across the complex. Its footprint is distributed almost evenly between the two antigens — roughly 840 Å^2^ on gH and 810 Å^2^ on gK — so that the epitope straddles the interprotein boundary rather than sitting on either partner alone (**Figure 5A and Figure 5C**). On gH, the epitope spans three discontinuous segments (residues ∼45–54, 136–142, and 253–277), and on gK it spans residues ∼191–201 and ∼280–308. These findings support our initial inference from the competition ELISA that group 1 mAbs recognize epitopes near the gH–gK interface (**Figure 3A**). Variations in the relative contributions of the two subunits to their epitopes could explain why group 1 contains both gH/gK-specific and gH/gL-cross-reactive mAbs despite their similar competition profiles. Superposition of B1L onto the HCMV gH/gL complex further suggests a structural basis for its gH/gK specificity, revealing a steric clash between the B1L CDRH3 and gL (**Figure S6B**).

In contrast to B1L, the footprint of D11L is heavily skewed toward gK: it buries ∼930 Å^2^ on gK versus only ∼380 Å^2^ on gH (**Figure 5A**). The epitope is centered on an extended, acidic stretch of gK, residues 197– 209, that lacks secondary structure and lies against the D11L heavy chain (**Figure 5A**). The interface of D11L-gH/gK is dominated by the heavy chain (**Figure 5B**), whereas the light chain makes only a peripheral contribution, contacting gH residues 35–39 and gK residues 206–209 (**Figure 5B**). Notably, whereas B1L and G2L bound in well-defined, rigid orientations, 3D classification revealed substantial conformational variability of D11L relative to gH/gK (**Figure S4**), likely arising from the flexibility of the gK segment spanning residues 191–211. This flexible mode of engagement is notable given the broad and strong FcγR activation mediated by D11L, suggesting a potential relationship between epitope flexibility and FcγR crosslinking and activation.

Together, our cryo-EM data define distinct mAb epitopes on HCMV gH/gK, providing a structural basis for gK-targeting by natural human antibodies.

### Immunization with gH/gK induces neutralizing and FcγR-activating antibodies in mice

Given the robust FcγR activation observed for gK-specific mAbs, we reasoned that immunization with the gH/gK protein complex could induce both gH-directed neutralizing antibodies and gK-specific antibodies capable of efficient FcγR activation. To test this, we assessed the immunogenicity of the gH/gK protein complex in mice.

To this end, BALB/c mice (n=5) were immunized with three doses of 30 µg gH/gK formulated 1:1 with AddaVax, and sera were analyzed at weeks 6 and 8 (2 and 4 weeks post-final dose, respectively) (**Figure 6A**). By week 8, anti-gH/gK IgG binding titers were readily induced, with a mean ED_50_ of approximately 9,300 (range of 4,343–18,911) (**Figure 6B and S7A**). Subclass profiling of gH/gK-targeting antibodies at week 8 showed a predominantly IgG1 response, with IgG1 being the major subclass in all mice, whereas mouse M2 also exhibited comparable IgG3 titers (**Figure 6C and S7B**).

**Figure 6.**
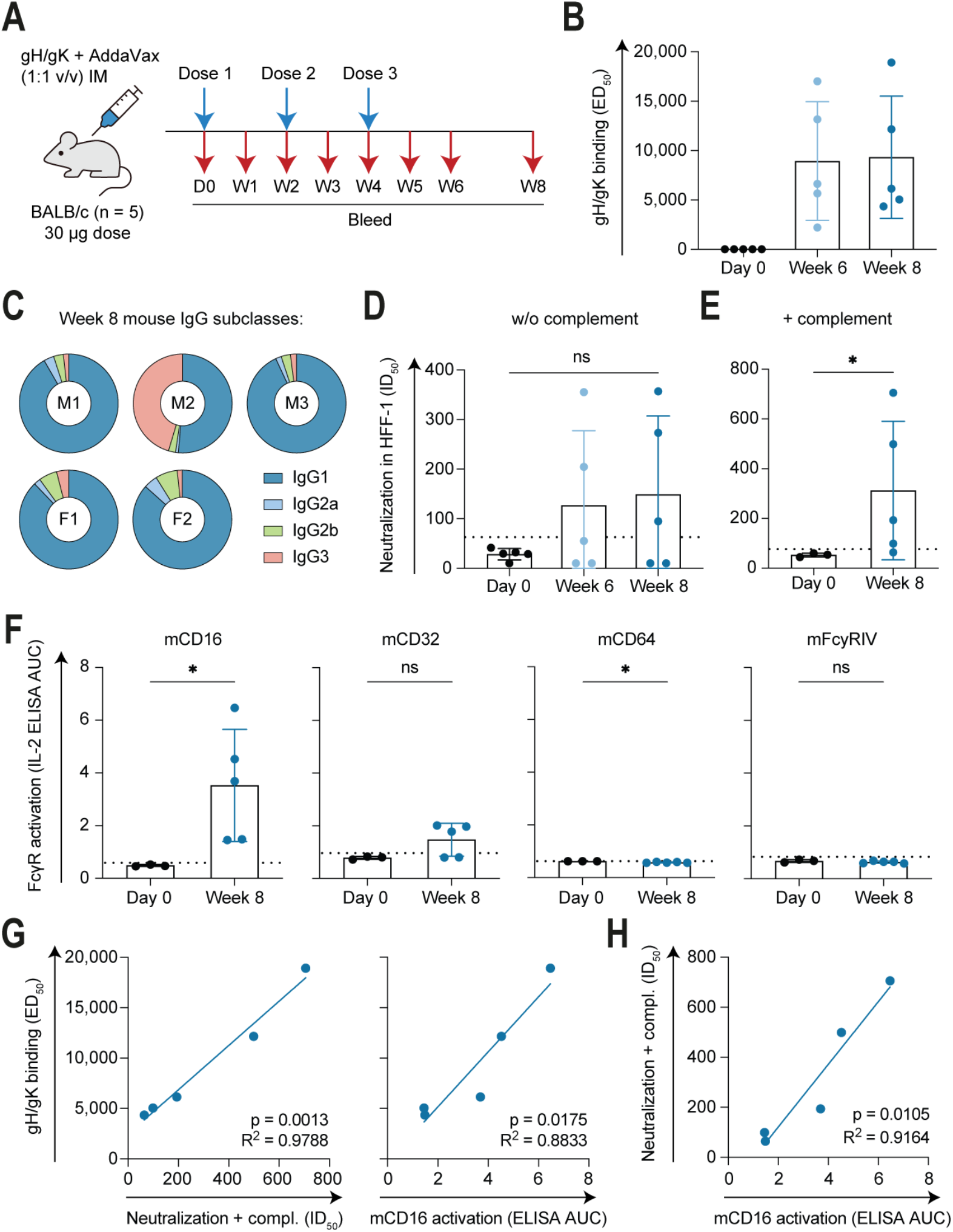
Immunization with gH/gK induces neutralizing and FcγR-activating antibodies in mice. (**A**) Mouse immunization strategy using the gH/gK complex. Five BALB/c mice were immunized intramuscularly with three doses of 30 µg gH/gK formulated 1:1 with AddaVax at 2-week intervals. Blood was collected on day 0 (D0) and weekly thereafter from week 1 to week 6 (W1–W6), with terminal bleeding at week 8 (W8). (**B**) gH/gK-complex binding titers, shown as ED_50_ values, in immunized mice at day 0, week 6, and week 8. ED_50_ values were calculated from serial dilution data shown in **Figure S7A**. Mean values and SDs are shown for each time point. (**C**) IgG subclass distribution of gH/gK-targeting antibodies in sera of immunized mice at week 8, shown for each mouse (M1–M3 and F1–F2). The color key for mouse IgG subclasses is shown below right. Percentages of each subclass were calculated from area under the curve (AUC) values derived from subclass-specific detection of gH/gK-reactive mouse serum IgG by ELISA, shown in **Figure S7B**. (**D–E**) Neutralizing activity of immunized mouse sera against HCMV_Towne_ infection of HFF-1 fibroblasts, shown as ID_50_ values and tested without complement (Figure 6D) or with complement (Figure 6E). Mouse sera were tested starting at a 1:10 dilution using an 8-point, 3-fold serial dilution series. The dotted lines indicate the positivity cutoff, calculated as the mean of day 0 samples plus 3 SD. Mean values and SDs are shown for each time point. Statistical significance was assessed using a Friedman test followed by Dunn’s multiple-comparisons test (Figure 6D) or a two-tailed Mann–Whitney test (Figure 6E). (**F**) Mouse FcγR activation by sera from immunized mice, measured for each receptor (mCD16, mCD32, mCD64, and mFcγRIV) using the BW reporter system under gH/gK complex coating conditions. Activation was measured by ELISA-based IL-2 detection in supernatants of the respective reporter cells. ELISA AUC values for day 0 and week 8 were calculated from the serial dilution data shown in **Figure S7D**. The dotted lines indicate the positivity cutoff, calculated as the mean of day 0 samples plus 3 SD. Mean values and SDs are shown for each time point. Statistical significance was assessed using a two-tailed Mann–Whitney test. (**G**) Left: correlation between gH/gK-complex binding titers (ED_50_ values; data from Figure 6B) and complement-enhanced HCMV_Towne_ neutralizing activity in HFF-1 fibroblasts (ID_50_ values; data from Figure 6E) in week 8 sera of immunized mice. Right: correlation between gH/gK-complex binding titers (ED50 values; data from Figure 6B) and mCD16 activation (ELISA AUC values; data from Figure 6F) in week 8 sera of immunized mice. Pearson correlation analysis was performed, and lines represent simple linear regression models. R^2^ and P values are indicated on each plot. (**H**) Correlation between complement-enhanced HCMV_Towne_ neutralizing activity in HFF-1 fibroblasts (ID_50_ values; data from Figure 6E) and mCD16 activation (ELISA AUC values; data from Figure 6F) in week 8 sera of immunized mice. Pearson correlation analysis was performed, and the line represents a simple linear regression model. R^2^ and P value are indicated on the plot.

HCMV-neutralizing activity of serum IgG from immunized animals, measured using HFF-1 fibroblasts, was detectable but lower relative to binding titers, with three of five mice exceeding the positivity cutoff (**Figure 6D**). From week 6 to week 8, neutralizing titers increased modestly on average (mean ID_50_ of 127 to 149) without reaching statistical significance compared with day 0 (**Figure 6D**). We also assessed neutralization of HCMV_Towne_ in fibroblasts in the presence of complement. Under these conditions, week 8 neutralizing titers were significantly higher on average compared with day 0, with a mean ID_50_ of 312 (**Figure 6E**).

We next assessed FcγR activation by serum IgG across different mouse Fcγ receptors in BW5147 reporter cell-based system^38^. First, to define the IgG subclass-specific activation potential, commercially available mouse IgG subclass control antibodies were tested for direct FcγR activation (**Figure S7C**). This analysis showed that mCD16 was activated by mIgG1, mIgG2a, and mIgG2b; mCD32 by mIgG1 only; mCD64 by mIgG2a only; and mFcγRIV by mIgG2a and mIgG2b (**Figure S7C**). Consistent with the predominantly mIgG1-subclass response induced by immunization, week 8 sera showed a significant increase in gH/gK-dependent mCD16 activation relative to day 0 (**Figure 6F and S7D**). A modest increase in mCD32 activation was also observed, with three mice exceeding the positivity cutoff (**Figure 6F and S7D**). In contrast, sera did not activate mCD64 or mFcγRIV, reflecting the insufficient induction of the required mIgG2a and mIgG2b subclasses by immunization (**Figure 6F and S7D**).

Importantly, gH/gK-binding IgG titers showed strong positive correlations with both HCMV neutralization (R^2^ = 0.9788, p = 0.0013) and mCD16 activation (R^2^ = 0.8833, p = 0.0175) (**Figure 6G**), indicating that these functions are largely titer-dependent. Consistent with these relationships, neutralization and mCD16 activation were also positively correlated (R^2^ = 0.9164, p = 0.0105), with mice demonstrating the highest neutralizing titers also exhibiting the strongest mCD16 activation (**Figure 6H**).

Collectively, these data show that gH/gK immunization elicits both HCMV-neutralizing and FcγR-activating antibodies in mice. Together with our human B cell and monoclonal antibody analyses, these findings highlight gH/gK as a promising vaccine candidate capable of engaging complementary antiviral antibody functions: pre-entry targeting of HCMV through neutralization and activation of FcγRs involved in the immune clearance of virions and infected cells.

## DISCUSSION

Neutralizing antibodies can efficiently block cell-free HCMV infection.^29–31^ Accumulating evidence further indicates a role for antibody responses beyond neutralization in limiting viral dissemination and congenital transmission, including through Fc-mediated effector functions.^32–35,42^ A protective role for antibodies is supported by gB/MF59 vaccine studies, in which IgG binding to gB was associated with a reduced risk of HCMV acquisition.^43^ Furthermore, passive administration of HCMV-specific hyperimmune globulin protected against placental transmission in a rhesus macaque model,^44^ while a neutralizing gH/gL-targeting mAb reduced fetal infection in a guinea pig model of congenital HCMV infection.^45^ Together, these observations suggest that an effective HCMV vaccine may benefit from eliciting both neutralizing and FcγR-engaging antibody responses.

Here, we identify gH/gK as a natural target of human humoral immunity. Across gH/gK-responsive donors, the concordance between plasma IgG binding and gH/gK-reactive memory B cell frequencies in different donors indicates that gH/gK reactivity is represented in both the circulating antibody and memory B cell compartments. These memory B cells are enriched for IgG3 in all gH/gK-seropositive donors. In parallel, repertoire features differed between donors, including distinct V_H_ gene usage. Together, these observations indicate the capacity for multi-epitope targeting and recruitment of diverse B cell precursors within the gH/gK-directed repertoire, which can be leveraged for immunogen design.

We show that B cell and antibody responses to gH/gK generate both neutralizing and FcγR-activating mAbs. Importantly, our data demonstrate that epitope position on gH/gK shapes antibody function. Our data support a model in which gH engagement is required for neutralization, whereas gK-specific antibodies are weakly or non-neutralizing but can act as potent activators in FcγR reporter assays, with broad FcγR activation exemplified by group 2 gH/gK-specific mAbs. Such antibodies, or their Fc-engineered variants,^35,46,47^ may provide a basis for future passive immunoprophylaxis or adjunctive therapeutic strategies for pregnant individuals and other high-risk groups, aimed at reducing transplacental transmission or limiting viral reactivation,^34,45^ pending rigorous evaluation in latency- and transmission-relevant models.

To our knowledge, we provide the first epitope-level analysis of FcγR activation by antibodies targeting HCMV entry glycoprotein complexes. Importantly, our data indicate that epitope context influences FcγR activation and that antibody binding alone is not sufficient for efficient Fc receptor activation. In our reporter assays using recombinant proteins, most mAbs were able to activate CD16aV but showed weak or absent activation of CD16aF and CD16b. Given the high prevalence of the CD16aF allele in human populations,^48^ this preferential activation of CD16aV highlights the potential value of antibodies that efficiently activate both CD16a variants. Notably, only mAbs from epitope group 2 activated both CD16a variants and CD16b to high levels in assays using recombinant gH/gK. Cryo-EM analysis of the representative group 2 mAb CS4p5_D11L revealed flexible engagement of its gK-specific epitope, with the Fab adopting multiple orientations relative to the gH/gK complex. This flexibility may reduce geometric constraints on Fc positioning and thereby facilitate productive FcγR engagement and crosslinking. Structure-guided selection of epitopes that promote Fab–Fc geometry, orientation, and spacing conducive to FcγR activation may therefore provide a promising path for HCMV immunogen design. Future mechanistic studies will be needed to define how epitope flexibility and antibody orientation regulate FcγR engagement.

Our virion-based assays further highlight the importance of antigen context and IgG subclass for FcγR activation. Cross-reactive group 4 mAbs that activated CD16aV with recombinant proteins induced minimal CD16 activation on virions, whereas stronger responses were observed for group 2 and 3 mAbs. For group 2 mAbs, virion-dependent FcγR activation was largely restricted to the IgG3 format. One possible explanation is that the extended IgG3 hinge provides additional freedom in the relative positioning of Fab and Fc domains, helping antibodies adapt to the spatial organization of antigens on the virion surface. These observations suggest that antibody geometry beyond the Fab–epitope interface may contribute to subclass-dependent FcγR activation, although the contribution of hinge flexibility in this setting remains to be established.

We also observed that gH/gK is relatively abundant on the surface of infected cells, consistent with reports that a substantial fraction of gH not incorporated into the gH/gL/gO trimer or gH/gL/UL128-131A pentamer is associated with gK on virions.^23^ This membrane abundance may favor opsonization and Fc clustering required for FcγR engagement.^49,50^ Defining the kinetics and tissue distribution of gH/gK expression during natural infection, latency, and reactivation will be important next steps.

Finally, in our mouse proof-of-concept immunization study, quantitative correlations between binding titers and both neutralization and mCD16 activation indicate that response magnitude is an important driver of these antiviral activities, supporting immunogen strategies that enhance the overall serum IgG response. The mouse IgG1-dominant response observed in our study preferentially mediated mCD16 and mCD32 activation, but not mCD64 or mFcγRIV, highlighting the need to tailor adjuvants and immunization regimens to broaden FcγR engagement. By analogy, human vaccine strategies may benefit from eliciting antibody subclasses that efficiently engage Fcγ receptors, particularly CD16a on NK cells.^51^

Beyond these humoral responses, cellular immunity plays an important role in controlling HCMV replication and preventing disease.^52,53^ HCMV-specific CD8^+^ and CD4^+^ T-cell responses are strongly associated with viral control, particularly in transplant recipients and congenital settings.^54–56^ Future studies should therefore determine whether gH/gK-based immunization elicits T-cell responses and how these might complement neutralizing and Fc-mediated antibody functions in providing protection.

In conclusion, our data position gH/gK as a vaccine candidate capable of coupling pre-entry targeting through viral neutralization with post-entry targeting through Fc-mediated effector mechanisms. These insights provide a blueprint for the development of an HCMV vaccine and antibody therapeutics that co-optimize neutralization and Fc functions, and motivate further evaluation of gH/gK-based strategies in infection-relevant models to assess their potential relevance to the prevention of congenital HCMV infection and transplantation-associated disease.

### Limitations of the study

Some aspects of this study should be considered when interpreting the findings. First, donor selection for B cell isolation was based on neutralization potency and breadth; therefore, the recovered repertoire may not fully represent the range of FcγR-activating responses present among all HCMV-seropositive individuals. Second, the memory B cell sort was not fully exclusive for gK because the gH/gL-negative gate was not maximally stringent, introducing some gH-reactive cells into the sorted population, although this also enabled characterization of epitopes spanning the gH/gK interface. Third, the mouse immunization experiments were performed with a small sample size using a single schedule, adjuvant, and protein platform, without head-to-head comparison to other gH-associated complexes; comparative studies and optimization of immunization strategy are needed to broaden neutralization and FcγR engagement. Finally, this study defines epitope-level structure-function relationships and does not assess efficacy; we did not test protection or quantify *in vivo* contributions of Fc-mediated functions to infection control. As HCMV is specific to human infection, designed challenge and maternal-fetal transmission models will be essential to link these correlates to outcomes.

## RESOURCE AVAILABILITY

### Lead contact

- Requests for further information and resources should be directed to and will be fulfilled by the lead contact, Matthias Zehner.

### Materials availability

- Requests for materials will be fulfilled by the lead contact upon request. Transfer of antibodies, protein constructs, and sample material might require a Material Transfer Agreement (MTA) for non-commercial usage.

### Data and code availability

- Antibody sequences generated in this study have been deposited in GenBank under accession numbers QB103584–QB103593.
- Cryo-EM maps have been deposited in the EMDB under accession codes EMD-79522 (global reconstruction), EMD-79524 (local reconstruction), and EMD-79526 (composite map). Atomic coordinates have been deposited in the PDB under accession code 39JB.
- All data that support the findings in this paper will be shared by the lead contact upon request.
- This paper does not report original code.
- Any additional information required to reanalyze the data reported in this paper is available from the lead contact upon request.

## ACKNOWLEDGMENTS

We gratefully thank all study participants for supporting our research by blood donation; all members of the Permar, Koch, Kolb, Hengel, Klein, Zehner, and McLellan Laboratories for helpful discussion and support. Moreover, we thank Anna Schmitt, Tina Bresser, and Lisa Kottege for lab management and assistance.

## AUTHOR CONTRIBUTIONS

Conceptualization, Artem A. and M.Z.; Methodology, Artem A., K.H., A.R.R., K.P.K., P.K., C.K., J.S.M., H.H., F.K., and M.Z.; Investigation, Artem A., K.H., Anna A., A.R.R., K.P.K., M.R., A.K.K., G.M.T., C.K., C.B., C.W., C.M.C., L.S., D.S., L.U., C.H., and M.K.; Visualization, Artem A., K.H., A.R.R., and K.P.K.; Formal analysis, Artem A., K.H., A.R.R., K.P.K., C.K., J.S.M., and M.Z.; Writing—original draft, Artem A., K.H., A.R.R., J.S.M., and M.Z.; Writing—review & editing, all authors; Funding acquisition, Artem A., M.K., J.S.M., H.H., F.K., and M.Z.; Project administration, Artem A., F.K., and M.Z.; Supervision, S.R.P., J.S.M., H.H., F.K., and M.Z.

## STAR★METHODS

### KEY RESOURCES TABLE

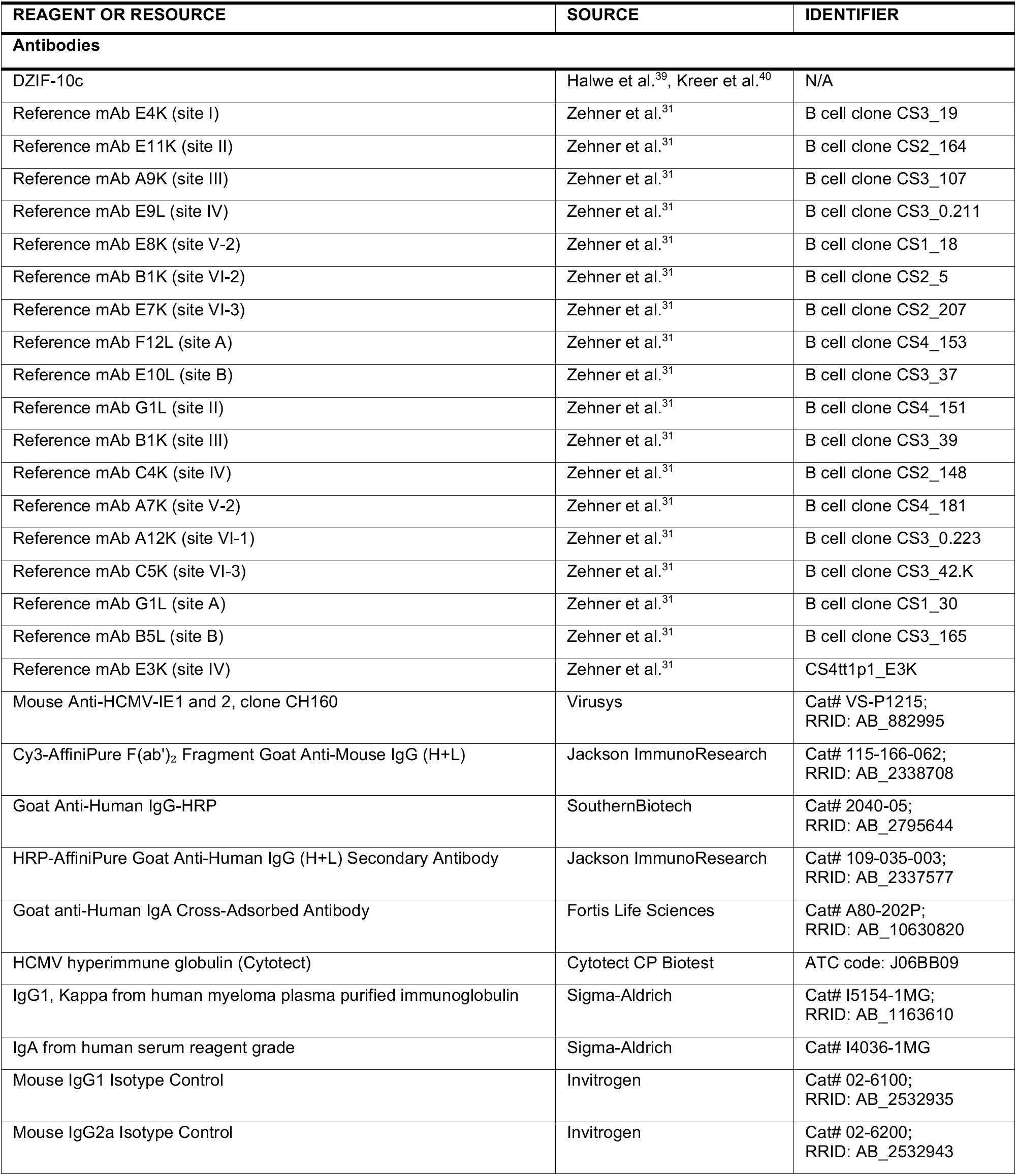
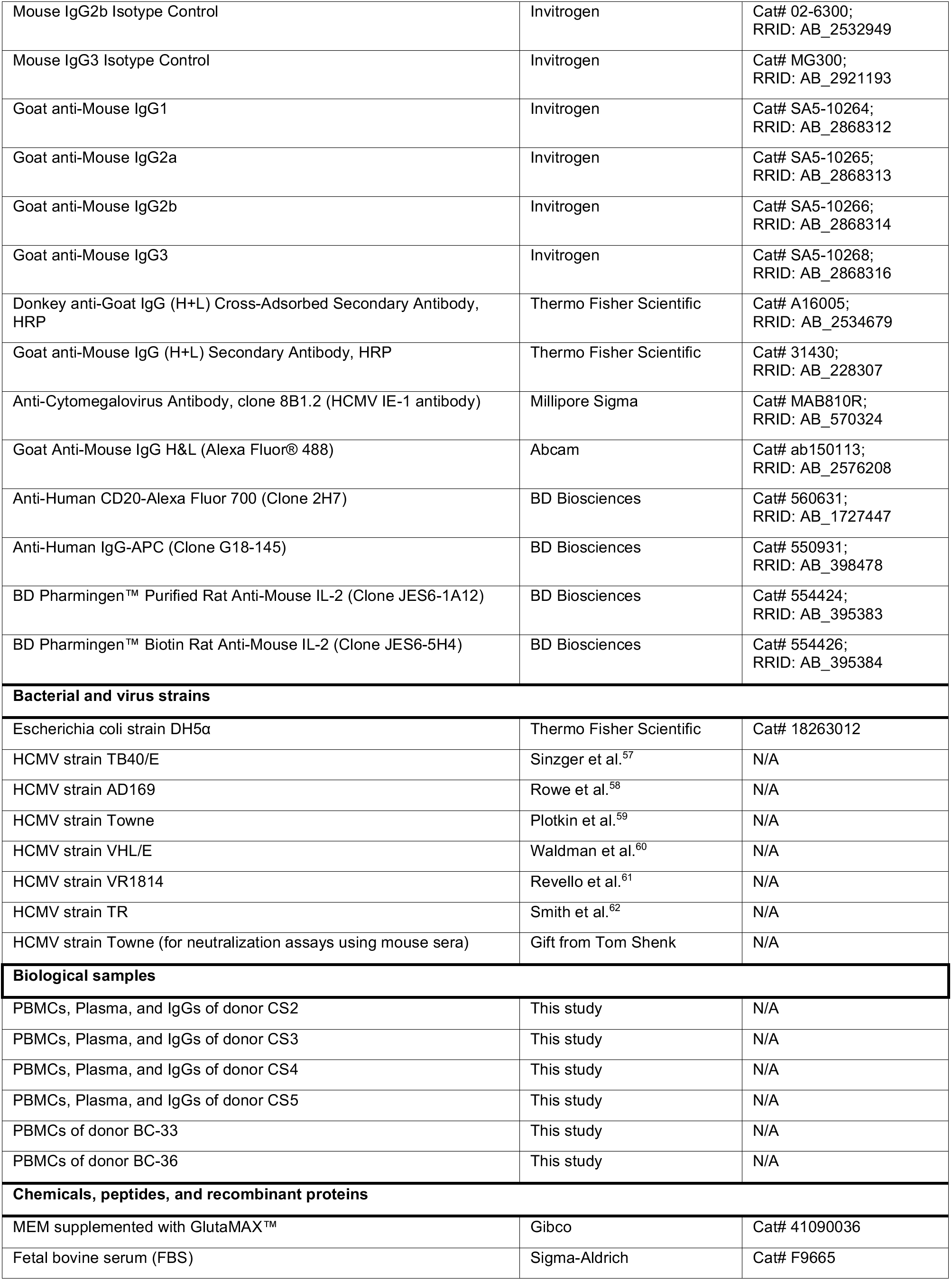
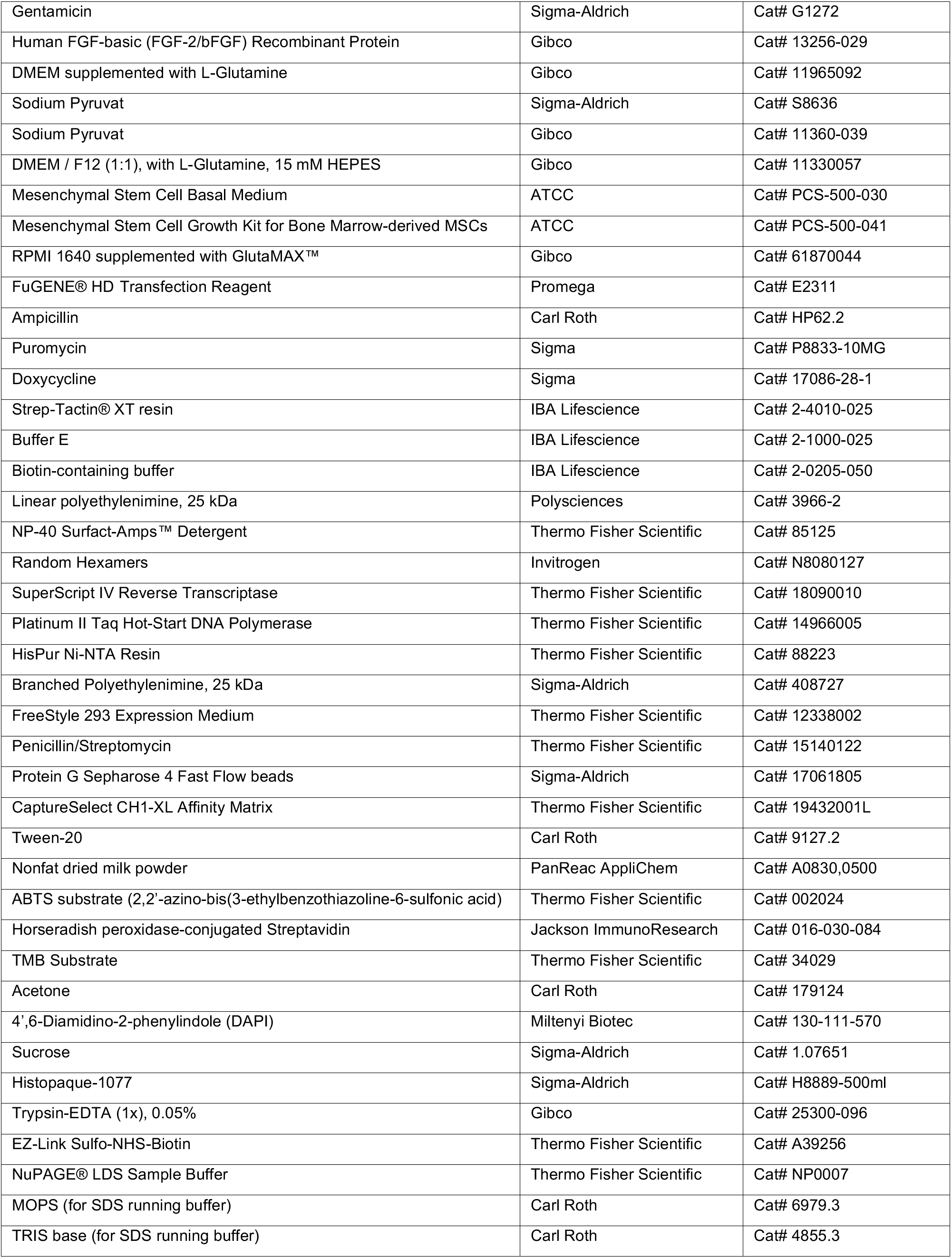
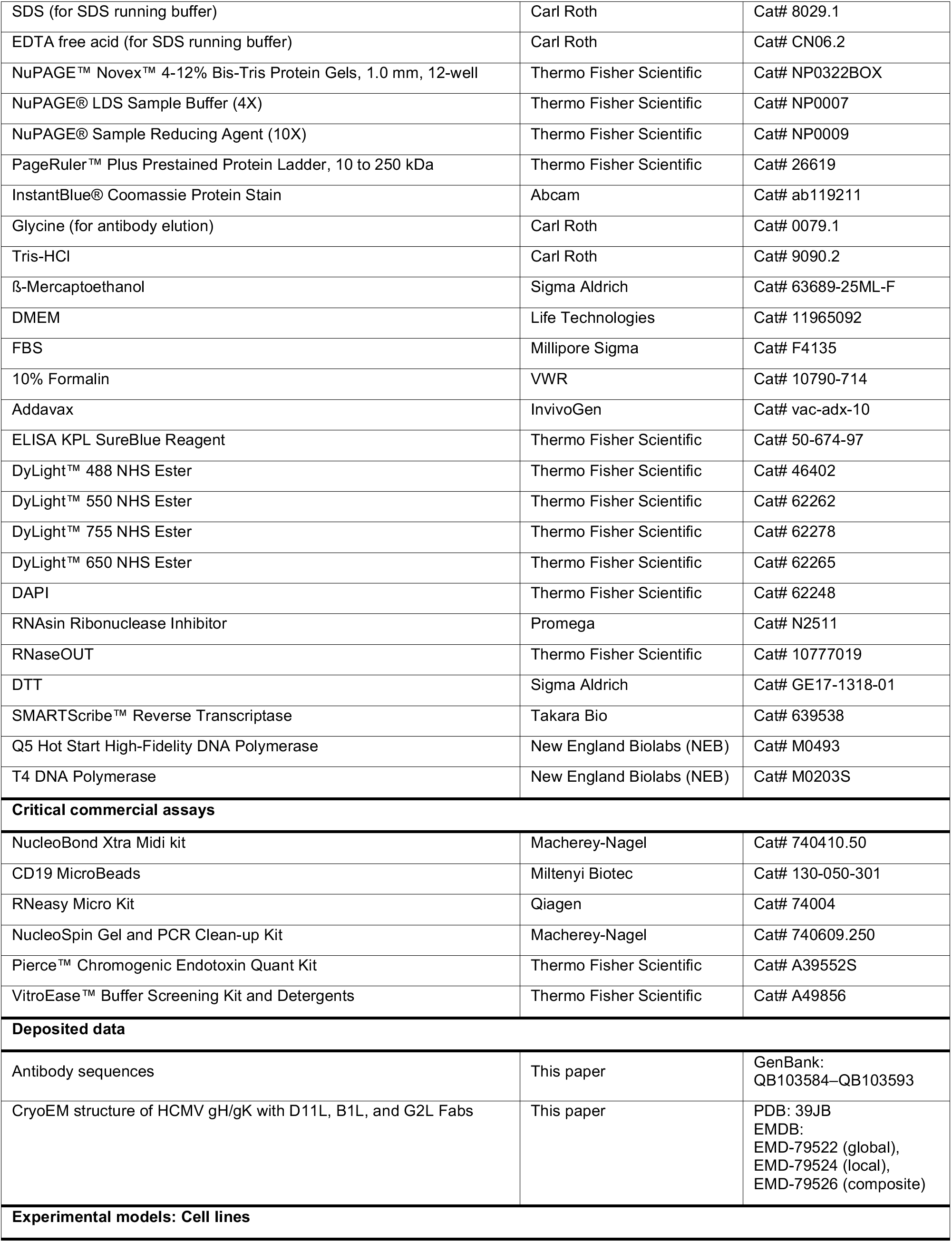
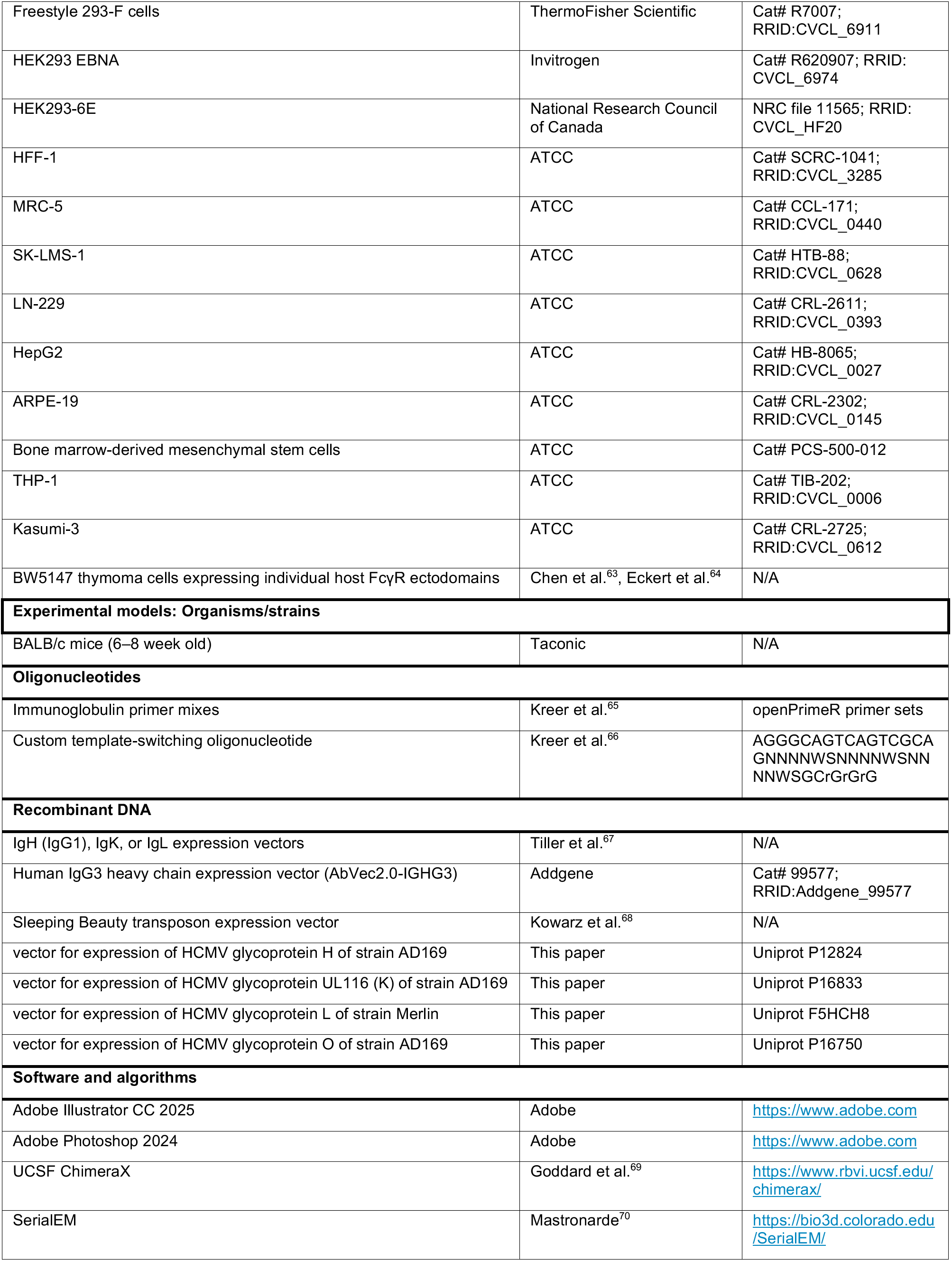
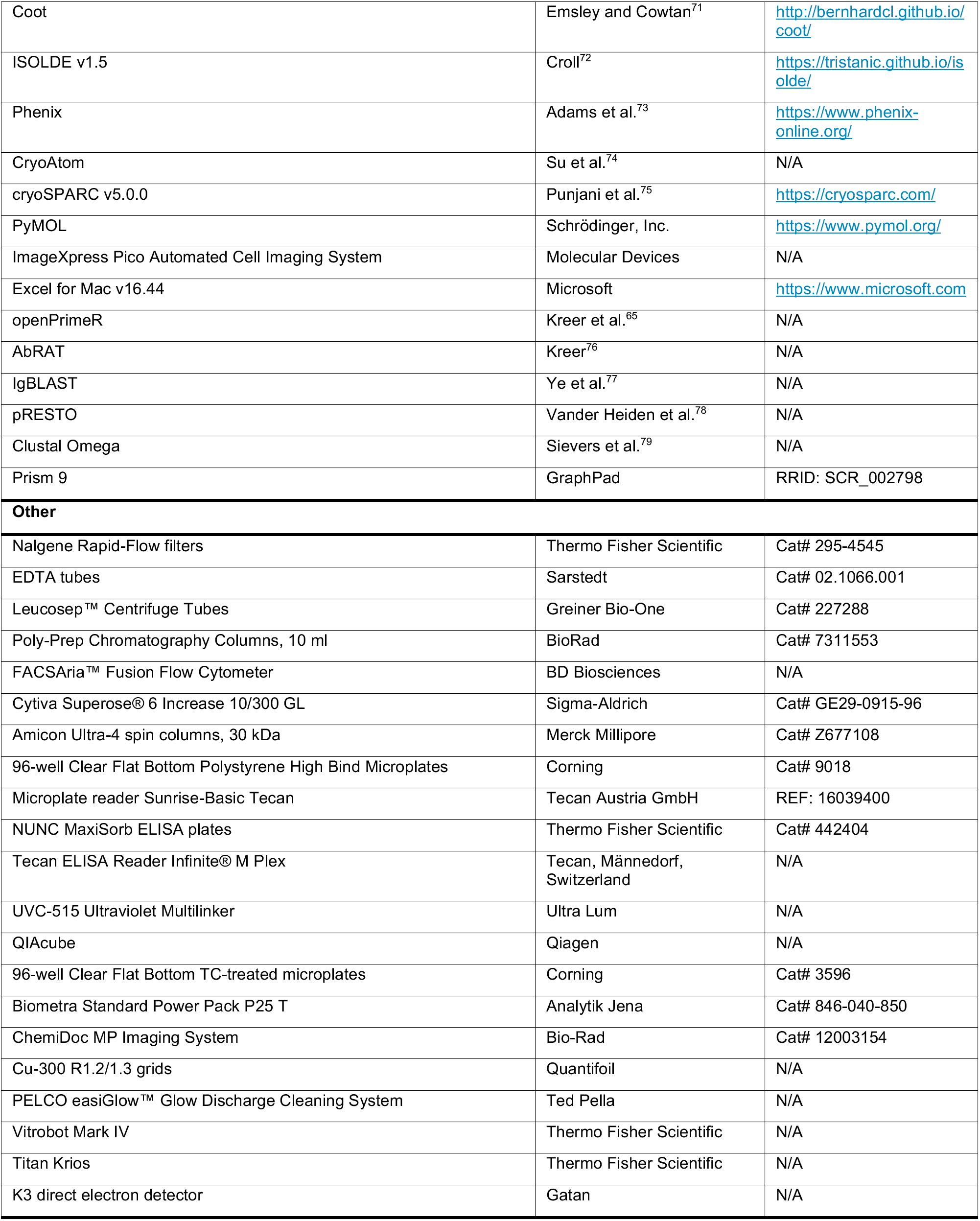

### EXPERIMENTAL MODEL AND STUDY PARTICIPANT DETAILS

#### HCMV-seropositive individuals and sample collection

The study included four HCMV-seropositive adult donors, designated CS2-CS5. Two participants were female (49 and 58 years old), and two participants were male (49 and 69 years old). Two HCMV-seronegative adult donors, designated BC-33 and BC-36, were additionally included as negative controls for the flow-cytometric analysis. No sex-stratified analysis was performed; information on gender, ancestry, race, and ethnicity was not available. The HCMV-seropositive participants were selected from a screening cohort of 9,000 HCMV-seropositive individuals.^36^ Plasma samples were ranked according to their capacity to neutralize the HCMV in human foreskin fibroblasts. 58 samples displayed exceptionally potent and broad neutralizing activity and were classified as elite neutralizers. From this group, four donors were enrolled for this study based on their response to the recall invitation, willingness to participate, and availability for blood collection. Blood samples were collected from donors who gave their written consent under a study protocol approved by the Institutional Review Board of the University of Cologne (study protocol #16-054).

#### Bacterial strains

Escherichia coli DH5α strain (Thermo Fisher Scientific) was used for molecular cloning and plasmid propagation. Bacteria were cultured in LB medium at 37°C, with 50 µg/mL ampicillin (Carl Roth) added for plasmid selection where appropriate.

#### Cells lines and primary cell cultures

HEK293-6E cells (National Research Council of Canada, NRC file 11565) were used for monoclonal antibody production and were cultured in FreeStyle 293 Expression Medium (Thermo Fisher) supplemented with 0.2% Penicillin-Streptomycin (Thermo Fisher) under continuous agitation at 37°C and 6% CO_2_. HEK293 EBNA cells (Invitrogen) were used to produce HCMV protein complexes and were cultured in FreeStyle 293 Expression Medium (Thermo Fisher) under continuous agitation at 37°C and 6% CO_2_. FreeStyle 293-F cells (Thermo Fisher) were used to produce the gH/gK complex for structural studies and were maintained in FreeStyle 293 Expression Medium (Gibco) under continuous agitation at 37°C and 8% CO_2_.

Cell lines and primary cultures used for HCMV infection and neutralization assays included HFF-1, MRC-5, SK-LMS-1, LN-229, HepG2, ARPE-19, bone marrow–derived mesenchymal stem cells (MSCs), THP-1, and Kasumi-3. These cells were maintained at 37°C and 5% CO_2_. HFF-1 fibroblasts (ATCC) were maintained in MEM supplemented with GlutaMAX™ (Gibco), 10% FBS (Sigma-Aldrich), 0.1 mg/mL gentamicin (Sigma-Aldrich) and 1 µg/L Human FGF-basic Recombinant Protein (Gibco). For neutralization assays using sera from immunized mice, HFF-1 cells were maintained in DMEM (Life Technologies) supplemented with 20% FBS (Millipore Sigma), 50 U/mL penicillin and 50 µg/mL streptomycin (Gibco). MRC-5, SK-LMS-1, LN-229 and HepG2 cell lines (ATCC) were maintained in DMEM supplemented with L-Glutamine (Gibco), sodium pyruvate (Sigma-Aldrich), 10% FBS and 0.1 mg/mL gentamicin. ARPE-19 cells (ATCC) were maintained in DMEM / F12 (1:1) supplemented with L-Glutamine, 15 mM HEPES (Gibco), 10% FBS and 0.1 mg/mL gentamicin. Bone marrow-derived MSCs (ATCC-PCS-500-012) were maintained in Mesenchymal Stem Cell Basal Medium (ATCC) supplemented with Mesenchymal Stem Cell Growth Kit for Bone Marrow-derived MSCs (ATCC) and 0.1 mg/mL gentamicin. THP-1 and Kasumi-3 cells (ATCC) were maintained in RPMI 1640 supplemented with GlutaMAX™ (Gibco), 10% FBS and 0.1 mg/mL gentamicin.

Mouse BW5147 thymoma cells (ATCC) were used to measure Fc gamma receptor activation by antibodies in BW reporter system.^38,80^ Thymoma cells and transfectants thereof were maintained in “RPMI BW medium”, containing RPMI 1640 supplemented with GlutaMAX™ (Gibco), 10% (v/v) heat-inactivated FBS, 1% (v/v) Penicillin-Streptomycin (Thermo Fisher), 1% (v/v) sodium pyruvate (100 mM, Gibco), and 0.1% (v/v) b-mercaptoethanol (Sigma-Aldrich). Cells were cultured at 37°C with 5% CO_2_ and split based on their growth rate. Cells were maintained at a density of 2×10^5^/ml to 1×10^6^/ml. Cells were tested regularly for mycoplasma contamination using PCR.

#### HCMV viruses

Replication-competent HCMV strains TB40/E, AD169, Toledo, Towne, VHL/E, VR1814, and TR were used in this study. To prepare stocks for mAb neutralization experiments, HFF-1 cultures were infected with the respective HCMV strains and incubated for 7 days. The resulting culture supernatants were transferred to fresh HFF-1 cells for a second 7-day propagation cycle. Supernatants collected after the second passage served as cell-free virus stocks and were aliquoted and stored at -80°C until use. Stock infectivity was quantified by fluorescence-based detection of HCMV immediate early antigens IE-1/2, as detailed in the “Virus neutralization assays” section.

For neutralization assays using sera from immunized mice, HCMV_Towne_ virus and bacterial artificial chromosomes (BACs) used for virus propagation were a gift from Professor Tom Shenk. HCMV_Towne_ was propagated on HFF-1 cells in T-175 culture flasks, and virus was collected from the supernatant and lysed infected cells when 100% of cells show cytopathic effects. Supernatant was cleared of cell debris by low-speed centrifugation, followed by ultracentrifugation through a 20% sucrose cushion. Virus titer was determined by incubating dilutions of virus stock in triplicate (starting 1:300, 6-point 3-fold dilutions) with HFF-1 cells for 72 hours at 37°C, followed by fixing with 10% formalin (VWR), staining with anti-HCMV IE-1 IgG (Millipore Sigma), and counter-staining with IgG-AF488 (Abcam) and 4’,6-diamidino-2-phenylindole (DAPI; Thermo Fisher Scientific). The titer was reported in PFU/mL.

#### Mouse Immunizations

All animal studies were conducted with approval by the Weill Cornell Medicine Institutional Animal Care and Use Committee (IACUC protocol: 2024-0013). 6–8 week old BALB/c mice (Taconic) were immunized thrice with 30 µg of immunogen (gH/gK), adjuvanted 1:1 (v/v) with Addavax (InvivoGen). Vaccine doses were stored on ice immediately after mixing and allowed to come to room temperature prior to administration. Antigens were administered by intramuscular injection (50µL) into the quadriceps. Animals were monitored 24 hours post immunization for adverse reactions. All mice were housed and fed according to institutional standard conditions (≤5 mice per cage, standard chow and water *ad libitum*). Blood was collected weekly by tail vein nick. Blood was incubated at room temperature for 30 minutes to enable clotting, followed by centrifugation for 6 minutes at 6000rpm (to separate serum from clotted whole blood). Serum was collected and stored at -80°C until use. At end of study mice were humanely euthanized by carbon dioxide asphyxiation and cervical dislocation, at which point blood was collected via cardiac puncture.

### METHOD DETAILS

#### Processing of blood from HCMV-seropositive donors and isolation of PBMCs and plasma IgG

Peripheral blood was collected into EDTA tubes (Sarstedt) and processed for the recovery of plasma and peripheral blood mononuclear cells (PBMCs). PBMCs were separated by density-gradient centrifugation using Leucosep tubes (Greiner Bio-One) containing Histopaque (Sigma-Aldrich), following the manufacturers’ instructions. The plasma fraction was retained separately and stored at -80°C.

For purification of total IgG, 1 mL of plasma was heat-inactivated at 56°C for 40 min and mixed with Protein G Sepharose (Sigma-Aldrich) at 4°C overnight. The resin–plasma mixture was subsequently loaded onto a 10 ml chromatography column (BioRad), and unbound material was removed by washing with PBS. Bound IgG was eluted with 0.1 M glycine (Carl Roth) at pH 3.0 and immediately neutralized with 1 M Tris-HCl (Carl Roth) at pH 8.0. The eluate was exchanged into PBS using 30 kDa Amicon centrifugal filters (Millipore). IgG concentrations were determined from absorbance at 280 nm using a NanoDrop One spectrophotometer, and purified samples were kept at 4°C until use.

#### HCMV glycoprotein production and purification

Recombinant HCMV glycoprotein complexes were produced using the Sleeping Beauty transposon system as previously described.^31^ For production of the gH/gK complex, two separate constructs included: gH (Uniprot P12824; HCMV strain AD169; AA 27–720; BM40 signal peptide; C-terminal Twin-Strep-tag) and gK (Uniprot P16833; HCMV strain AD169; AA 24–314; BM40 signal peptide). For production of the gH/gL complex, two separate constructs included: gH (Uniprot P12824; HCMV strain AD169; AA 27–720; BM40 signal peptide; C-terminal Twin-Strep-tag) and gL (Uniprot F5HCH8; HCMV strain Merlin; AA 32–278; BM40 signal peptide). For production of the gH/gL/gO complex, three separate constructs included: gH (Uniprot P12824; HCMV strain AD169; AA 27–720; BM40 signal peptide; C-terminal Twin-Strep-tag), gL (Uniprot F5HCH8; HCMV strain Merlin; AA 32–278; BM40 signal peptide), and gO (Uniprot P16750; HCMV strain AD169; AA 40–466; BM40 signal peptide). The corresponding expression plasmids were transfected into HEK293 EBNA cells (RRID: CVCL_6974) together with the transposase plasmid at a 10:1 ratio using FuGENE® HD transfection reagent (Promega GmbH, Madison, USA). Stable transfectants were selected with 2 µg/mL puromycin (Sigma-Aldrich) and expanded in triple-layer culture flasks. Glycoprotein expression was induced by adding 0.5 µg/mL doxycycline (Sigma-Aldrich). Culture supernatants were collected from confluent cells, filtered, and subjected to affinity purification using Strep-Tactin® XT resin (IBA Lifescience, Göttingen, Germany). Bound protein complexes were eluted using a biotin-containing buffer (IBA Lifesciences), dialyzed against PBS, aliquoted, and stored at -80°C. Before use in the mouse immunization study, endotoxin levels in the purified gH/gK complex were measured using the Pierce™ Chromogenic Endotoxin Quant Kit (Thermo Fisher Scientific) according to the manufacturer’s instructions and were ≤0.031 EU per 30 µg immunization dose.

For structural analyses, the gH construct comprises an immunoglobulin signal peptide, ectodomain residues 24–718 of HCMV gH strain AD169 (UniProt P12824), followed by a hexa-histidine tag. The gK construct comprises ectodomain residues 1–314 of HCMV UL116 strain AD169 (UniProt P16833), followed by a Twin-Strep-tag. The constructs were separately cloned into the mammalian expression vector pcDNA3.1(+). Plasmids encoding gH and gK were transiently co-transfected at a 1:2 gH:gK DNA ratio into FreeStyle™ 293-F (Thermo Fisher Scientific; RRID:CVCL_6911) cultures using PEI (Polysciences). 5 μM kifunensine was added approximately 4 hours after transfection. After 6–7 days, medium was harvested by centrifugation, 0.22 µm filtered, then passed over Strep-Tactin Sepharose resin (IBA Lifesciences) by gravity, washed with 3 column volumes of 1× PBS, and eluted with 1× Buffer E (100 mM Tris-HCl pH 8.0, 150 mM NaCl, 1 mM EDTA, and 2.5 mM desthiobiotin) (IBA Lifesciences). The elution was concentrated with Amicon Ultra centrifugal filters (Millipore), and flash-frozen in liquid nitrogen. Samples were rapidly thawed and hard spun just before injection onto a Superose 6 Increase 10/300 GL column (Cytiva). The SEC running buffer was composed of 2 mM Tris pH 8.0, 200 mM NaCl, and 0.02% (w/v) sodium azide. Main peak fractions were pooled, concentrated, and flash-frozen with liquid nitrogen.

#### SDS-PAGE analysis of HCMV glycoprotein complexes

For SDS-PAGE analysis, purified recombinant HCMV glycoprotein complexes were lysed in NuPAGE® LDS Sample Buffer (Thermo Fisher Scientific), denatured at 70°C for 10 min, and separated by electrophoresis in MOPS/SDS running buffer in NuPAGE™ Novex™ 4-12% Bis-Tris Protein Gels (Thermo Fisher Scientific). For reducing SDS-PAGE, NuPAGE® Sample Reducing Agent (Thermo Fisher Scientific) was additionally used. After electrophoresis, gel was stained for 30 min with InstantBlue® Coomassie Protein Stain (Abcam). After destaining with deionized water, colorimetric signal was detected in a ChemiDoc MP Imaging System (Bio-Rad).

#### Depletion ELISA for determination of plasma IgG residual binding activities

For determination of residual binding activities after depletion, 96-well High Bind microplates (Corning) were coated overnight at 4°C with the respective target protein at 3 µg/mL in PBS. Plates were washed three times with PBST (PBS containing 0.05% Tween 20; Carl Roth) and blocked for 1 h at room temperature (RT) with 5% nonfat dry milk (PanReac AppliChem) in PBST. Purified total IgG was analyzed in 5-fold serial dilutions starting at 2,000 µg/mL. Before application to the plates, IgG samples were incubated overnight at 4°C with the respective depletion protein at a 15-fold excess of total IgG to occupy antibodies recognizing the depletion antigen (e.g. antibodies directed against gH/gL epitopes). The resulting mixtures were transferred to the antigen-coated plates and incubated for 90 min at RT. After washing, bound human IgG was detected using horseradish peroxidase-conjugated goat anti-human IgG (SouthernBiotech, 2040-05; 1:2000 in PBST with 5% milk powder). Plates were incubated for 45 min at RT and washed before addition of 2,2′-azino-bis(3-ethylbenzothiazoline-6-sulfonic acid) substrate (ABTS; Thermo Fisher Scientific). Absorbance was measured at 415 nm with reference subtraction at 695 nm using an absorbance plate reader (Tecan) after a fixed development time. Binding curves were generated and analyzed in GraphPad Prism 9.

#### Isolation of HCMV-specific and whole repertoire IgG+ B cells

B cells were enriched from PBMC preparations using CD19 MicroBeads (Miltenyi Biotec) according to the manufacturer’s instructions. For flow cytometric sorting, two preparations of the HCMV gH/gK complex were independently conjugated to DyLight 488 and DyLight 550 (Thermo Fisher Scientific), respectively, and gH/gL was conjugated to DyLight 755 (Thermo Fisher Scientific) according to the manufacturer’s instructions. CD19-enriched cells were stained with the fluorescently labeled glycoprotein complexes, DAPI (Thermo Fisher Scientific), anti-human CD20-Alexa Fluor 700 (BD Biosciences), and anti-human IgG-APC (BD Biosciences). DAPI^−^CD20^+^IgG^+^ cells negative for gH/gL and positive for both gH/gK probes were sorted as single cells using a FACSAria™ Fusion Flow Cytometer (BD Biosciences) into 96-well plates. Each well contained 4 µL of FACS buffer comprising 0.5x PBS, 0.5 U/µL RNasin (Promega), 0.5 U/µL RNaseOUT (Thermo Fisher Scientific), and 10 mM dithiothreitol (DTT; Sigma Aldrich). Plates were stored at -80°C until further processing.

For total B cell repertoire sequencing, CD19-enriched cells were stained with DAPI, anti-human CD20-Alexa Fluor 700, and anti-human IgG-APC. DAPI^−^CD20^+^IgG^+^ cells from each donor were sorted into tubes containing FBS (Sigma Aldrich). After sorting, cells were collected by centrifugation at 400g for 5 min, the supernatants were then removed, and the cell pellets were stored at -80°C until further processing.

#### Amplification and analysis of B cell receptor sequences

B cell receptor heavy and light chain sequences were amplified from single sorted B cells using a previously described workflow.^81^ After thawing the plates on ice, single cells were lysed by adding a mixture containing NP-40 (Thermo Fisher Scientific), random hexamers (Invitrogen), and RNaseOUT (Thermo Fisher Scientific). Plates were incubated at 65°C for 5 min and subsequently cooled on ice. The released RNA was reverse-transcribed using SuperScript IV Reverse Transcriptase (Thermo Fisher Scientific) in the presence of RNaseOUT and RNasin (Promega). Heavy and light chain variable regions were subsequently amplified from the resulting cDNA by sequential semi-nested PCR using optimized V gene-specific primer mixes (openPrimeR^65^), Platinum Taq Hot Start DNA Polymerase (Thermo Fisher Scientific), and 6% KB Extender. Amplicons of the expected size were identified by gel electrophoresis and subjected to Sanger sequencing.

Raw Sanger sequencing data were processed and analyzed using the Antibody Repertoire Analysis Toolkit (AbRAT).^76^ For sequence quality control (QC), nucleotide calls with Phred scores below 16 were masked. Sequences passed QC if they had a mean Phred score of at least 28, contained fewer than 15 internal masked nucleotides, and were at least 240 nucleotides long. Sequences were annotated using IgBLAST^77^ and trimmed to encompass the variable region starting from framework region 1 (FWR1) through the end of the J gene. Only productive sequences that passed QC were retained for repertoire and clonal analyses.

Clonal relationships were determined separately for each donor in AbRAT using the iterative CDRH3-similarity clustering algorithm. In detail, heavy chain sequences were grouped according to their V gene and J gene assignments and clustered based on CDRH3 amino acid similarity using a length threshold of 1, a normalized Levenshtein-distance threshold of 0.25, and 20 iterations. Thus, sequences assigned to the same V and J genes, with a CDRH3 length difference of no more than one amino acid and at least 75% CDRH3 amino acid sequence similarity, were considered clonally related. All QC-filtered B cell receptor sequences were included in analyses of IgG subclass distribution, CDRH3 length, and V_H_ gene germline identity. For the analysis of V_H_ gene usage, donor CS3 was excluded because of the low number of recovered B cells, sequences from all other donors were included.

#### Next generation sequencing of IgG-positive B cell repertoires

Total B-cell receptor repertoires were generated using template-switching reverse transcription PCR followed by Illumina Miseq sequencing, as described previously.^66^ In brief, RNA was extracted from frozen cell pellets using the RNeasy Micro Kit (Qiagen) on a QIAcube instrument (Qiagen) according to the manufacturer’s instructions. 10 µL aliquot of each RNA eluate was used for cDNA synthesis with SMARTScribe Reverse Transcriptase (Takara), following the SMARTer RACE 5′/3′ protocol. Reverse transcription used a custom template-switching oligonucleotide containing an 18-nucleotide unique molecular identifier (UMI). Heavy, κ, and λ chain variable regions were amplified by semi-nested PCR using constant region-specific primers and Q5 High-Fidelity DNA Polymerase (New England Biolabs). Amplicons were separated by gel electrophoresis and recovered using the NucleoSpin Gel and PCR Clean-up Kit (Macherey-Nagel). Purified heavy, κ, and λ chain amplicons from each donor were combined at a 2:1:1 ratio. Library preparation and paired-end sequencing on an Illumina MiSeq platform with a read length of 2x300 bp were performed at the Cologne Center for Genomics. Raw reads were filtered for a mean Phred score of at least 25 and a minimum length of 250 bp. Sequence processing and assembly were performed using the pRESTO toolkit,^78^ and immunoglobulin gene annotation was performed using IgBLAST^77^. Reads sharing the same UMI were aligned with Clustal Omega^79^ for error correction and consensus-sequence generation. Only productive consensus sequences derived from at least three paired-end reads sharing the same UMI and showing complete alignment to the corresponding V gene reference sequence were included in downstream repertoire analyses.

#### Cloning and in vitro production of monoclonal antibodies

Monoclonal antibodies were generated using a previously described workflow^82^ with modifications. In short, heavy and light chain variable region sequences were obtained as eBlocks Gene Fragments (IDT) containing terminal overlapping sequences required for cloning. The fragments were inserted into the corresponding human IgH (IgG1), IgK, or IgL expression vectors^67^ by sequence- and ligation-independent cloning (SLIC)^83,84^ using T4 DNA Polymerase (NEB). For production of IgG3 variants, the corresponding heavy chain variable regions were cloned into the human IgG3 expression vector AbVec2.0-IGHG3 (Addgene #99577). The resulting plasmids were propagated in chemically competent E. coli DH5α (Thermo Fisher Scientific) and verified by Sanger sequencing. Sequence-verified heavy and light chain plasmids were co-transfected into HEK293-6E cells (National Research Council Canada; RRID: CVCL_HF20) for antibody expression using branched polyethylenimine (PEI, Sigma-Aldrich). 7 days post-transfection, supernatants were collected and incubated with Protein G Sepharose 4 Fast Flow beads (Sigma-Aldrich) overnight at 4°C. The beads with bound antibodies were subsequently loaded onto a 10 ml chromatography columns (BioRad), and unbound material was removed by washing with PBS. Bound antibodies were eluted with 0.1 M glycine (Carl Roth) at pH 3.0 and immediately neutralized with 1 M Tris-HCl (Carl Roth) at pH 8.0. The eluate was exchanged into PBS using 30 kDa Amicon centrifugal filters (Millipore). Antibody concentrations were determined from absorbance at 280 nm using a NanoDrop One spectrophotometer, and purified samples were kept at 4°C until use.

#### ELISA for monoclonal antibody binding activity

For binding ELISA, 96-well High Bind microplates (Corning) were coated overnight at 4 °C with the respective target protein at 3 µg/mL in PBS. Plates were washed three times with PBST (PBS containing 0.05% Tween 20; Carl Roth) and blocked for 1 h at room temperature (RT) with 5% nonfat dry milk (PanReac AppliChem) in PBST. Monoclonal antibodies were analyzed in 5-fold serial dilutions starting at 30 µg/mL and were transferred to the antigen-coated plates and incubated for 90 min at RT. After washing, bound antibodies were detected using horseradish peroxidase-conjugated goat anti-human IgG (SouthernBiotech, 2040-05; 1:2000 in PBST with 5% milk powder). Plates were incubated for 45 min at RT and washed before addition of 2,2′-azino-bis(3-ethylbenzothiazoline-6-sulfonic acid) substrate (ABTS; Thermo Fisher Scientific). Absorbance was measured at 415 nm with reference subtraction at 695 nm using an absorbance plate reader (Tecan) after a fixed development time. Binding curves and EC_50_ values were generated and analyzed using GraphPad Prism 9. Positive binding was defined by an increasing absorbance with a final signal of at least 0.4 (gH/gK) or 0.2 (gH/gL) and an EC50 < 150 µg/mL.

#### Antibody-competition ELISA

Prior to the competition ELISA, each antibody was biotinylated using the EZ-Link Sulfo-NHS-Biotin (Thermo Fisher) according to the manufacturer’s instructions, and a buffer exchange to PBS was performed using 30 kDa Amicon centrifugal filters (Millipore). For antibody competition ELISA, 96-well High Bind microplates (Corning) were coated overnight at 4°C with 10 µg/mL of non-biotinylated antibodies (1^st^ mAb) in PBS. Plates were washed three times with PBST (PBS containing 0.05% Tween 20; Carl Roth) and blocked for 1 h at room temperature (RT) with 5% nonfat dry milk (PanReac AppliChem) in PBST. Next, gH/gK complex at a concentration of 3 µg/mL in 5% nonfat dry milk in PBST was added and incubated with the coated antibodies for 90 minutes at RT. After another three washes, biotinylated antibodies were added at 10 µg/mL concentration and incubated for 90 minutes at RT. Then, plates were incubated with horseradish peroxidase-conjugated Streptavidin (Jackson ImmunoResearch, 1:5000 in PBST / 5% Nonfat dried milk powder) for 30 min at RT and washed before addition of 2,2′-azino-bis(3-ethylbenzothiazoline-6-sulfonic acid) substrate (ABTS; Thermo Fisher Scientific). Absorbance was measured at 415 nm with reference subtraction at 695 nm using an absorbance plate reader (Tecan) after a fixed development time. Optical density values were sequentially normalized for each biotinylated antibody and unlabeled antibody and converted to percent competition, with 0% representing uninhibited binding of the biotinylated antibody and 100% representing maximal competition. Competition heatmaps were generated and analyzed using GraphPad Prism 9.

#### HCMV neutralization assays with monoclonal antibodies

Neutralizing activity of mAbs against HCMV infection was evaluated in HFF-1, MRC-5, SK-LMS-1, LN-229, HepG2, ARPE-19, bone marrow–derived mesenchymal stem cells (MSCs), THP-1, and Kasumi-3 cells. Assays were performed using the cell type-specific media described above; for HFF-1 and MSCs neutralization assays, MEM supplemented with GlutaMAX™ (Gibco), 10% FBS (Sigma-Aldrich), and 0.1 mg/mL gentamicin (Sigma-Aldrich) was used.

For assays in HFF-1, MRC-5, SK-LMS-1, LN-229, HepG2, ARPE-19, or MSCs cells, 96-well Clear Flat Bottom TC-treated microplates (Corning) were seeded with 15,000 cells per well and incubated overnight at 37°C. THP-1 and Kasumi-3 cells were seeded at 50,000 cells per well in RPMI 1640 supplemented with GlutaMAX™ and 0.1 mg/mL gentamicin without FBS, subsequent neutralization steps were performed in the complete RPMI 1640 medium containing 10% FBS.

Antibodies were serially diluted in the corresponding cell type-specific medium. For neutralization assays in different host cell lines, mAbs were tested at a starting concentration of 10 µg/mL. For neutralization of different HCMV strains in HFF-1 cells, mAbs were tested at a starting concentration of 50 µg/mL. Antibody dilutions were mixed at a 1:1 ratio with the indicated HCMV strain at the multiplicity of infection (MOI) of 1 infectious unit per cell or below (MOI of approximately 0.5–1.0). Virus–antibody mixtures were incubated for 2 h at 37°C before addition to the cells. Following incubation for 18–20 h at 37°C, cells were fixed with 80% acetone (Carl Roth) for 5 min at room temperature and washed three times with PBST (PBS containing 0.05% Tween 20; Carl Roth). Next, plates were stained with mouse mAb targeting HCMV immediate early antigen 1/2 (IE 1/2; 1:1000 in PBS; clone CH160; Virusys; VS-P1215) overnight at 37°C. After three washes with PBST, plates were incubated for 1 h at 37°C with Cy3-AffiniPure F(ab’)2 Fragment Goat Anti-Mouse IgG (H+L) (Jackson ImmunoResearch; RRID: AB_2338708) diluted 1:300 in PBS and 1 µg/mL DAPI (Miltenyi Biotec), followed by three additional washes.

Fluorescence images were acquired using an ImageXpress Pico system. DAPI-stained nuclei were used to identify and enumerate cells, and infection was quantified as the number of IE 1/2-positive cells relative to the total number of cells. Infection levels were expressed relative to the no-antibody virus-only control, which was defined as 100% infection. Dose–response curves were fitted by nonlinear regression using GraphPad Prism 9 to determine neutralizing capacity (%) and potency (IC_50_ values, µg/mL) of each antibody. Neutralizing capacity was defined as the difference between 100% infection (virus-only control) and the lower plateau (minimum) of the dose-response curve. Thus, a lower plateau of 30% residual infection corresponded to a neutralizing capacity of 70%. If no clear lower plateau was reached, residual infection at the highest antibody concentration tested was used to calculate the maximal observed neutralizing capacity. IC_50_ values were calculated only for antibodies with a neutralizing capacity of at least 50%, and correspond to the antibody concentration at which 50% of the maximal neutralization effect was achieved (relative IC_50_).^85^

#### Surface staining of HCMV-infected fibroblasts

Before use in the assay, monoclonal antibodies were conjugated to DyLight 650 (Thermo Fisher Scientific), according to the manufacturer’s instructions. For surface staining, 96-well Clear Flat Bottom TC-treated microplates (Corning) were seeded with 15,000 HFF-1 cells/well in MEM supplemented with GlutaMAX™ (Gibco), 10% FBS (Sigma-Aldrich) and 0.1 mg/mL gentamicin (Sigma-Aldrich) and incubated overnight at 37°C. Next, cells were infected with TB40/E strain of HCMV at the multiplicity of infection (MOI) of 0.5 infectious units per cell and incubated for 5 days at 37°C. Staining of HCMV proteins expressed on the cell surface was performed before fixation of the cells to limit intracellular staining. First, non-HCMV specific monoclonal human antibody (DZIF-10c^39,40^) at a concentration of 500 µg/mL was added for 1 h at 37°C to block HCMV-encoded Fc receptors expressed on surface of infected cells. Cells were then washed manually with PBS by carefully dispensing liquid against the side of each well to minimize cell detachment. Next, DyLight 650–conjugated HCMV-specific antibodies at a concentration of 4 µg/mL were incubated with infected cells for 1 h at 37°C. After manual washing with PBS, cells were fixed with 80% acetone (Carl Roth) for 5 min at room temperature and washed three times with PBST (PBS containing 0.05% Tween 20; Carl Roth). Plates were stained with 1 µg/mL DAPI (Miltenyi Biotec) for 10 min at 37°C and washed three times with PBST. Fluorescence images covering the entire well were acquired in the DAPI and DyLight 650 channels using an ImageXpress Pico system. Cells were identified based on DAPI-stained nuclei, and the average DyLight 650 pixel intensity was quantified across all cells in each well.

#### FcγR activation assays

The assay was performed as described earlier.^38^ For antigen-independent direct activation assays and assays using recombinant glycoprotein complexes, ELISA MaxiSorp plates (Thermo Fisher Scientific) containing antibodies, either coated directly on the assay plates or bound to immobilized antigen (gH/gK or gH/gL/gO at 2 µg/mL coated overnight at 4 °C) were co-cultured with BW5147 reporter cells (50,000/well in 200 µL media) expressing individual host FcγR ectodomains^63,64^ for ∼16 h at 37°C in a 5% CO_2_ atmosphere. Reporter cell mIL-2 secretion, correlating to receptor activation, was quantified by subsequent anti-mIL-2 sandwich ELISA.

For virion-based assays, 20,000 virions of HCMV_TB40/E_ strain were suspended in 50 µL PBS and added to 96-well Clear Flat Bottom TC-treated microplates (Corning), followed by incubation overnight at 4°C. Immobilized virions were subsequently inactivated by UV-crosslinking at 3700kJ twice, using the UVC-515 Ultraviolet Multilinker. After two washes with PBST, wells were blocked with 300 µL/well of blocking buffer for at least 1 hour at room temperature. Following a single PBST wash, gH/gK-targeting antibodies (50 µg/mL) were added in 50 µL PBS per well and incubated overnight at 4°C. Subsequently, wells were washed three times with RPMI 1640, and BW5147 reporter cells were added at 50,000 cells/well in 200 µL media. Plates were then incubated for 24 hours at 37°C with 5% CO_2_. Reporter cell mIL-2 secretion, correlating to receptor activation, was quantified by subsequent anti-mIL-2 sandwich ELISA.

The level of mIL-2 secreted upon activation of BW5147 reporter cells was measured in anti-mIL-2 sandwich ELISA as described earlier.^38,86^ Briefly, ELISA MaxiSorp plates (Thermo Fisher Scientific) were coated with 1 µg/mL of rat anti-mouse-IL-2 antibody (BD Biosciences, clone JES6-1A12, #554424) in PBS and incubated overnight at 4°C. Plates were washed and blocked. Supernatants from the BW5147-FcγR reporter assay were transferred to the mIL-2-ELISA plates. Supernatants were incubated for overnight at 4°C on the ELISA plate, and wells were subsequently washed five times. 1 µg/mL of biotinylated rat anti-mouse-IL-2 (BD Biosciences, clone JES6-5H4, #554426) in ELISA blocking buffer was added and incubated for 90 minutes at RT. Plates were washed five times, and Streptavidin-Peroxidase (Jackson Immuno Research, #016-030-084) 1:1000 in blocking buffer was added for 30 minutes at RT. Wells were washed five times, and 50 µL of ELISA TMB 1-Step™ Ultra substrate solution (Thermo Fisher Scientific) was added, followed by 50 µL of 1 M H_2_SO_4_ to stop the reaction. Absorbance was measured using a Tecan ELISA Reader Infinite® M Plex (Tecan, Männedorf, Switzerland) at a wavelength of 450 nm and a reference wavelength of 620 nm.

To verify that tested antibodies remained on the plate during incubation with the reporter cells, the original assay plates, still containing either antigen-bound or directly immobilized antibodies, were further analyzed after removal of the mIL-2-containing supernatants. The respective secondary antibody (1:3000 goat anti-human IgG-HRP or goat anti-human IgA-HRP) was added, and the resulting signal was designated “residual binding.”

FcγR activation and residual binding data were reported as optical density (OD) values. When both readouts were assessed, the results were presented as paired activation and residual binding heatmaps generated and analyzed using GraphPad Prism 9.

For dose–response analyses, serial dilutions of antibodies were applied to gH/gK-coated plates, and all subsequent assay steps were performed as described above. CD16aV activation curves were fitted using a variable-slope four-parameter logistic nonlinear regression model in GraphPad Prism 9.

For analysis of direct mouse FcγR activation, commercially available recombinant mouse IgG1, IgG2a, IgG2b, and IgG3 subclass control antibodies were coated on ELISA plates at a starting concentration of 1 µg/mL, followed by 2-fold serial dilutions. All subsequent assay steps were performed as described above using BW5147 reporter cells expressing mCD16, mCD32, mCD64, or mFcγRIV.^63^ FcγR activation was reported as mIL-2 sandwich ELISA OD values.

For analysis of mouse sera, day 0 and week 8 serum samples from gH/gK-immunized mice were applied to gH/gK-coated plates at a starting dilution of 1:1000, followed by 2-fold serial dilutions. All subsequent assay steps were performed as described above using BW5147 reporter cells expressing mCD16, mCD32, mCD64, or mFcγRIV. AUC values were calculated from the mIL-2 sandwich ELISA OD values across the serial dilution series with the baseline set to zero using GraphPad Prism 9.

#### Expression and purification of Fabs for structural studies

A stop codon was introduced before the hinge region of heavy chains to generate antigen-binding fragments (Fabs). FreeStyle™ 293-F (Thermo Fisher Scientific) cultures were transiently co-transfected with plasmids encoding heavy and light chains using PEI. After 6–7 days, medium was harvested by centrifugation, 0.22 µm filtered, then passed over CaptureSelect™ IgG-CH1 Affinity Matrix (Thermo Fisher Scientific). Bound Fab was eluted with 100 mM glycine pH 3.0. The elution was immediately neutralized with 1 M Tris pH 8.0 and then concentrated and buffer exchanged into SEC running buffer using Amicon Ultra centrifugal filter units (Millipore Sigma). Fabs were stored at 4°C, or flash-frozen with liquid nitrogen for long-term storage.

#### Cryo-EM sample prep and data collection

Purified HCMV gH/gK was rapidly thawed, hard spun, and diluted to a final concentration of 3.5 mg/mL in buffer containing 2 mM Tris pH 8.0, 200 mM NaCl, and 0.02% (w/v) sodium azide. Each Fab (B1L, D11L, and G2L) was added in a 1.4:1 Fab:gH/gK ratio. The combined complex was allowed to incubate at room temperature for 30 minutes. Immediately before freezing, 0.5 µL of β-OG (Octyl-β-D-Maltoside) (VitroEase™ Buffer Screening Kit and Detergents, Thermo Fisher Scientific) was mixed with 3.5 µL of the complex, bringing the final detergent concentration to 0.25× CMC. 4 µL of sample was applied to Cu-300 R1.2/1.3 grids (Quantifoil) that had been glow discharged for 30 seconds at 20 mA and 0.39 mBar (PELCO easiGlow™, Ted Pella). Using a Vitrobot Mark IV (Thermo Fisher Scientific), a blot force of 1 was applied for 10 seconds to blot away excess liquid before plunge-freezing into liquid ethane. Samples were blotted in 100% humidity at 22°C. 4,000 movies were collected from a single grid at 105,000× magnification corresponding to a calibrated pixel size of 0.8332 Å on an FEI Titan Krios operating at 300 kV and equipped with a K3 direct electron detector (Gatan). All movies were collected using SerialEM v4.0.10 automation software.^70^ Additional details about data collection parameters can be found in **Table S1**.

#### Cryo-EM processing and structure building

Motion correction, CTF estimation, particle picking, and 2D classification were performed using CryoSPARC v5.0.0 live processing.^75^ A preliminary volume was generated using 63,877 particles and further refined with C2 symmetry. It appeared that the addition of β-OG detergent influenced the formation of dimer complexes. The C2 volume was used to generate templates, and template picking was performed on 3,804 movies. Two rounds of 2D classification were performed: first with 80 classes and an uncertainty factor of 1, followed by 384,921 selected particles further sorted into 80 classes with an uncertainty factor of 3. 193,919 particles were selected and a subset of 100,000 particles were used to perform an ab initio reconstruction with four classes followed by heterogeneous refinement of those four classes and the C2 volume using all selected particles. A monomer complex class was identified and non–uniform refined to a final resolution of 3.27 Å. As the D11L Fab was poorly resolved and blurry in this global refinement map, 3D classification was performed using a focus mask to sort the particles by D11L position. The 3D classes revealed a flexible epitope, with some discrete states likely stabilized by the neighboring B1L Fab. The class that appeared to be most stabilized was further refined and used for local refinement to resolve the D11L epitope. A composite map of this local refinement and the global refinement was generated in Phenix v1.21.^73^ Initial models were generated by CryoAtom v2.1^74^ and the structure was iteratively refined using a combination of PyMOL v2.5.2 (Schrodinger), Coot,^71^ Phenix v1.21,^73^ ChimeraX v1.12,^69^ and ISOLDE v1.12.^72^ Map validations and the cryo-EM processing workflow can be found in **Figure S4 and Figure S5**.

#### Mouse serum IgG binding assay

Serum IgG binding to the immunogen gH/gK was assessed at weeks 0, 6, and 8 using ELISA at a starting sera dilution of 1:1000, followed by 8-point, 3-fold serial dilutions. 384-well plates were coated with 1 µg/mL antigen overnight at 4°C, followed by blocking in assay diluent (1x PBS [pH 7.4] containing 4% whey, 15% normal goat serum, and 0.5% Tween 20). IgG binding was detected by peroxidase-conjugated goat anti-mouse IgG secondary antibody (Thermo Fisher Scientific) and developed using SureBlue substrate (Thermo Fisher Scientific). Optical densities (OD) were obtained using a Synergy plate reader. Data was reported as the 50% effective dose (ED_50_) using non-linear regression analysis on GraphPad Prism 9.

#### ELISA for mouse IgG subclass determination

For IgG subclass determination, ELISA plates were coated overnight at 4°C with 2 µg/mL gH/gK in PBS. After washing and blocking, week 8 serum samples from immunized mice were tested in 2-fold serial dilutions ranging from 1:1000 to 1:64000 and incubated with coated gH/gK overnight at 4°C. Bound antibodies were detected using subclass-specific goat anti-mouse IgG1, IgG2a, IgG2b, or IgG3 antibodies at 0.5 µg/mL, followed by HRP-conjugated donkey anti-goat IgG at 0.3 µg/mL. ELISA TMB 1-Step™ Ultra substrate solution (Thermo Fisher Scientific) was used for development. Absorbance was measured using a Tecan ELISA Reader Infinite® M Plex (Tecan, Männedorf, Switzerland) at a wavelength of 450 nm and a reference wavelength of 620 nm.

Subclass-specific binding was reported as optical density (OD) values. AUC values were calculated from the serial dilution data with the baseline set to zero using GraphPad Prism 9. For each mouse, the relative contribution of each IgG subclass was calculated by dividing its AUC by the sum of the IgG1, IgG2a, IgG2b, and IgG3 AUCs.

#### Mouse serum IgG neutralization assay

Sera samples were heat inactivated at 56°C for 30 minutes prior to assays. Fibroblast neutralization of HCMV_Towne_ was assessed on HFF-1 cells; 6,000 cells were seeded per well of a 384-well plate and incubated overnight. HCMV_Towne_ at an MOI of 1.5 was incubated with sera IgG at a 1:10 starting sera dilution followed by 8-point 3-fold serial dilutions. Rabbit complement was added at a 1:8 dilution, when used. Virus-antibody complexes were incubated with cells for 24 hours, followed by fixing with 10% formalin (VWR), staining for HCMV IE-1 (Millipore Sigma) and counter-staining with IgG-AF488 (Abcam) and 4’,6-diamidino-2-phenylindole (DAPI; Thermo Fisher Scientific). Images were acquired on the ImageXpress Pico Automated Cell Imaging System (Molecular Devices). Infected cells were calculated as a percentage of AF488-positive cells relative to the total number of cells. Data was reported as the 50% infectious dose (ID_50_) and calculated using non-linear regression analysis (Sigmoidal 4-parametric logistic) on GraphPad Prism, with average of virus infection set as the maximum constraint for data and zero as the minimum constraint.

### QUANTIFICATION AND STATISTICAL ANALYSIS

All statistical details for individual experiments can be found in corresponding figure legends, results and method details sections. Statistical significance was defined as follows: ns, p ≥ 0.05; *p < 0.05; **p < 0.01; ***p < 0.001; and ****p < 0.0001. Quantitative and statistical analyses were performed using GraphPad Prism 9 and Microsoft Excel for Mac (v16.112). Microscopic quantification and analyses were done by ImageXpress Pico microscope. For structural analyses cryoSPARC, Phenix, Coot, CryoAtom, ISOLDE, UCSF ChimeraX and PyMOL were used.

