## Supplementary Materials for "B cell response to HCMV gH/UL116 elicits neutralizing and FcγR-activating antibodies"

### **SUPPLEMENTAL INFORMATION**

#### **Supplementary Figures**

**Figure S1.** Characterization of gH/gK-reactive plasma IgG and memory B cell responses, Related to Figure 1.

**Figure S2.** Additional characterization of reference mAbs and group 4 cross-reactive mAbs, Related to Figure 3.

**Figure S3.** Additional characterization of FcγR activation by gH/gK-targeting and reference mAbs, Related to Figure 4.

**Figure S4.** Cryo-EM data processing workflow, Related to Figure 5.

**Figure S5.** Cryo-EM map validation, Related to Figure 5.

**Figure S6.** HCMV gH/gK vs GATE and Trimer, Related to Figure 5.

**Figure S7.** Additional characterization of gH/gK-reactive antibody responses and FcγR activation in immunized mice, Related to Figure 6.

#### **Supplementary Table**

**Table S1.** Cryo-electron microscopy data collection and refinement statistics, Related to STAR Methods.

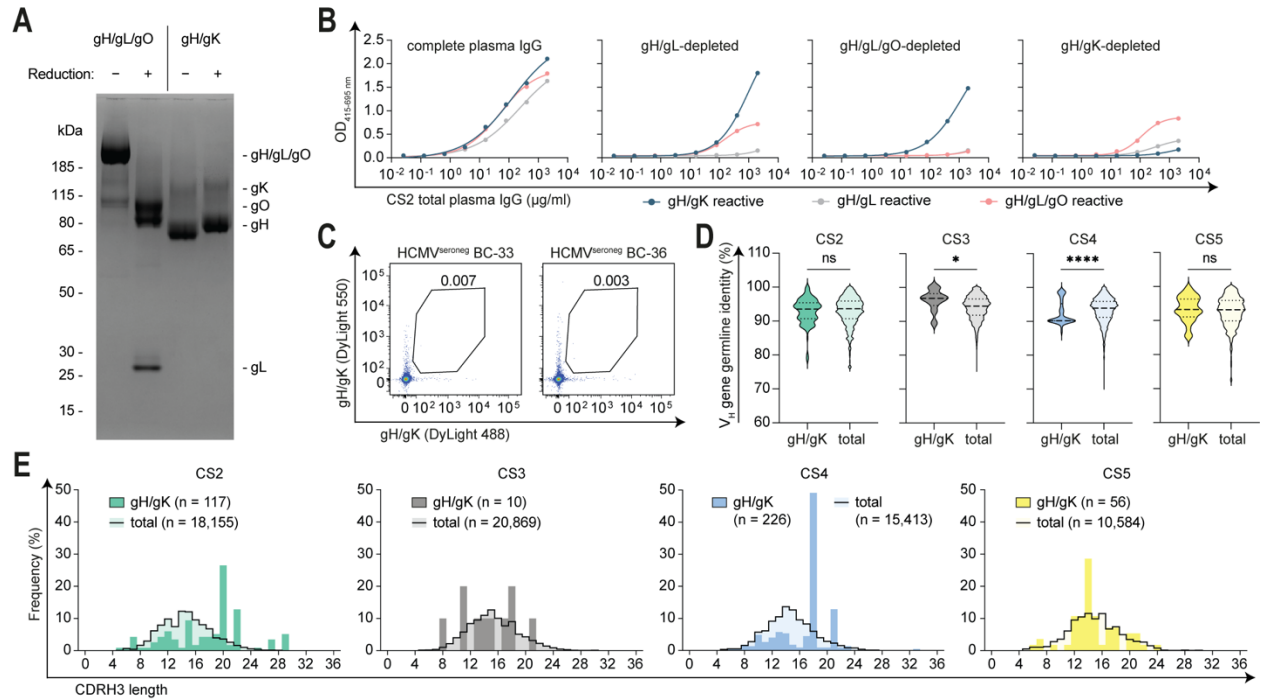

**Figure S1. Characterization of gH/gK-reactive plasma IgG and memory B cell responses.**

(A) Non-reducing (–) and reducing (+) SDS-PAGE of gH/gL/gO and gH/gK protein complexes, stained with Coomassie brilliant blue. (B) ELISA analysis of CS2 total plasma IgG reactivity against different HCMV protein complexes. Reactivity is shown for total plasma IgG (first graph), gH/gL-depleted plasma IgG (second graph), gH/gL/gO-depleted plasma IgG (third graph), and gH/gK-depleted plasma IgG (fourth graph). Reactivity against gH/gK, gH/gL, and gH/gL/gO is shown in blue, gray, and pink, respectively. Representative data of at least two separate experiments is shown. OD, optical density. (C) FACS analysis of memory B cells from HCMV-seronegative donors BC-33 and BC-36, with the percentage of gH/gK<sup>+</sup> cells indicated in each plot. (D) Violin plots showing V<sub>H</sub> gene DNA germline identity of isolated B cell receptor sequences from gH/gK-reactive B cells from each donor compared with the corresponding whole IgG-positive B cell repertoires, determined by NGS. Medians are indicated by dashed lines and quartiles by dotted lines. Significance was assessed using the Kolmogorov–Smirnov test. (E) Distribution of CDRH3 amino acid lengths among gH/gK-reactive B cells from each donor compared with the corresponding whole B cell repertoires determined by NGS. Colored bars indicate the frequency of gH/gK-reactive B cells at each CDRH3 length, and black lines indicate the corresponding whole IgG-repertoire distributions.

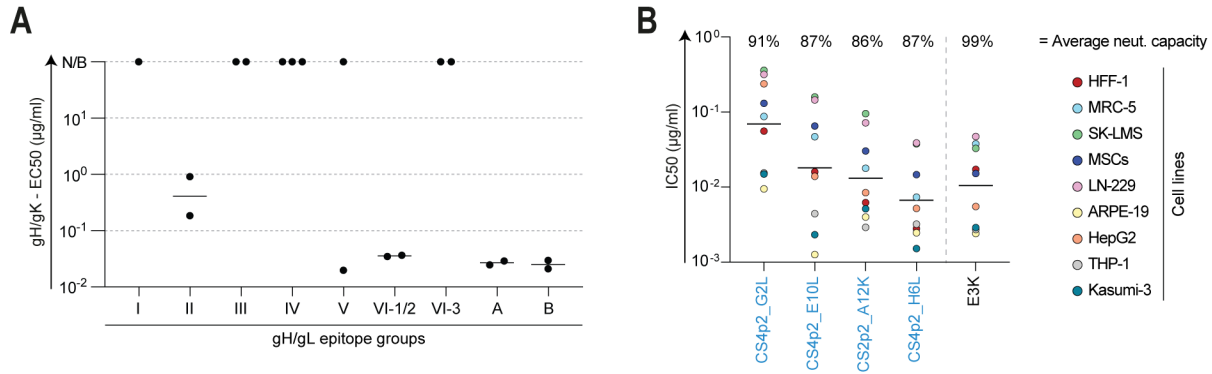

**Figure S2. Additional characterization of reference mAbs and group 4 cross-reactive mAbs.**

(A) Binding activity of 18 mAbs from the gH/gL-targeting reference panel<sup>31,37</sup> to the gH/gK complex, shown as ELISA-derived EC<sub>50</sub> values (μg/ml). mAbs are grouped according to gH/gL epitope groups. Each dot represents one mAb and shows the geometric mean of at least two separate experiments. Black lines indicate geometric means of all mAbs within the same epitope group. N/B, non-binding. (B) Neutralizing activity of group 4 cross-reactive mAbs against HCMV<sub>TB40/E</sub> infection in different host cell lines, shown as IC<sub>50</sub> values (μg/ml), compared with the previously described gH/gL-specific broadly neutralizing mAb CS4tt1p1\_E3K (referred to as E3K). IC<sub>50</sub> values for cross-reactive mAbs are derived from **Figure 3D** and were calculated from two separate experiments. The average neutralizing capacity across all cell lines for each mAb is shown above. Neutralizing capacity was calculated from mAb dose-response curves and represents the difference between the infection rate in no-antibody conditions and the residual infection plateau at saturating mAb concentrations (also see Methods).

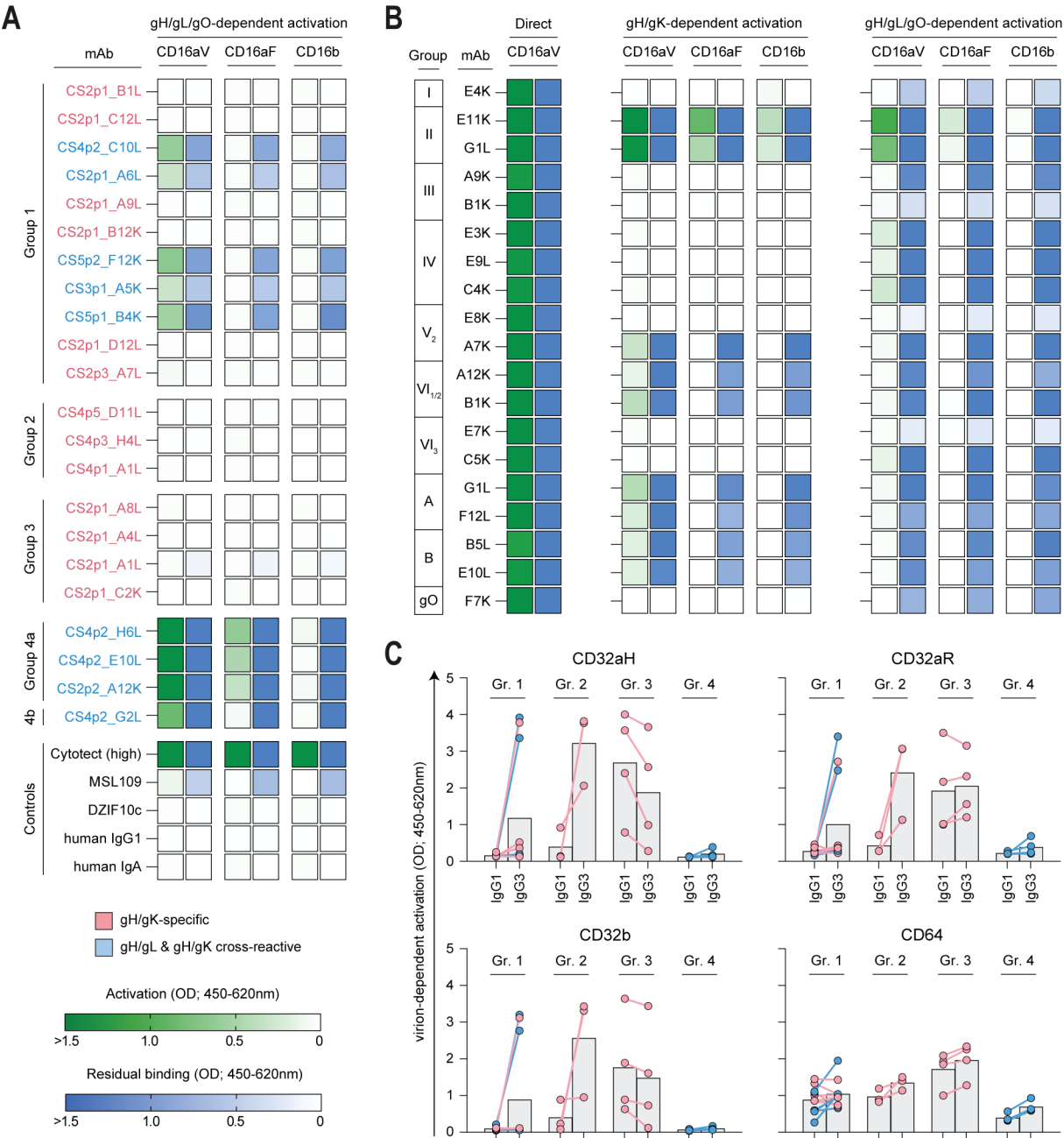

(Caption on the next page.)

**Figure S3. Additional characterization of FcγR activation by gH/gK-targeting and reference mAbs.**

**(A)** Human CD16 activation by gH/gK-reactive mAbs measured using the BW5147 reporter cell system under gH/gL/gO complex coating conditions. For each mAb, condition, and CD16 variant (CD16aV, CD16aF, and CD16b), paired heatmap squares show CD16 activation in green and residual antibody binding in blue. Activation was measured by ELISA-based mIL-2 detection in supernatants of the respective reporter cells; residual binding of antibodies remaining after each activation readout was measured by ELISA within the same assay. gH/gK-specific mAbs are indicated by pink text labels, and cross-reactive mAbs by blue text labels. mAbs are arranged by epitope group, with control antibodies shown below. mAbs were used at 1 µg/ml, Cytotect was used at 10 µg/ml. Each antibody was tested in two separate experiments; representative data from one experiment are shown. **(B)** Human CD16 activation by reference mAbs measured using the BW5147 reporter cell system under antigen-independent mAb coating conditions for direct activation (left), gH/gK complex coating conditions (center), and gH/gL/gO complex coating conditions (right). For each mAb, condition, and CD16 variant (CD16aV, CD16aF, and CD16b), paired heatmap squares show CD16 activation in green and residual antibody binding in blue. Activation was measured by ELISA-based mIL-2 detection in supernatants of the respective reporter cells; residual binding of antibodies remaining after each activation readout was measured by ELISA within the same assay. mAbs are arranged according to gH/gL epitope groups and were used at 1 µg/ml. **(C)** Human FcγR activation by gH/gK-reactive mAbs, measured using BW5147 reporter cells expressing CD32aH, CD32aR, CD32b, or CD64 under virion-dependent conditions with immobilized HCMV<sub>TB40/E</sub> virions. Activation was measured by ELISA-based IL-2 detection in supernatants of the respective reporter cells. Dots show activation by individual mAbs, with lines connecting the IgG1 and IgG3 measurements for the same mAb. Pink and blue dots indicate gH/gK-specific and gH/gL-cross-reactive mAbs, respectively. mAbs are arranged by epitope group, and gray bars indicate mean activation for each group and IgG subclass. mAbs were used at 50 µg/ml. Each antibody was tested in at least three separate experiments; representative data from one experiment are shown.

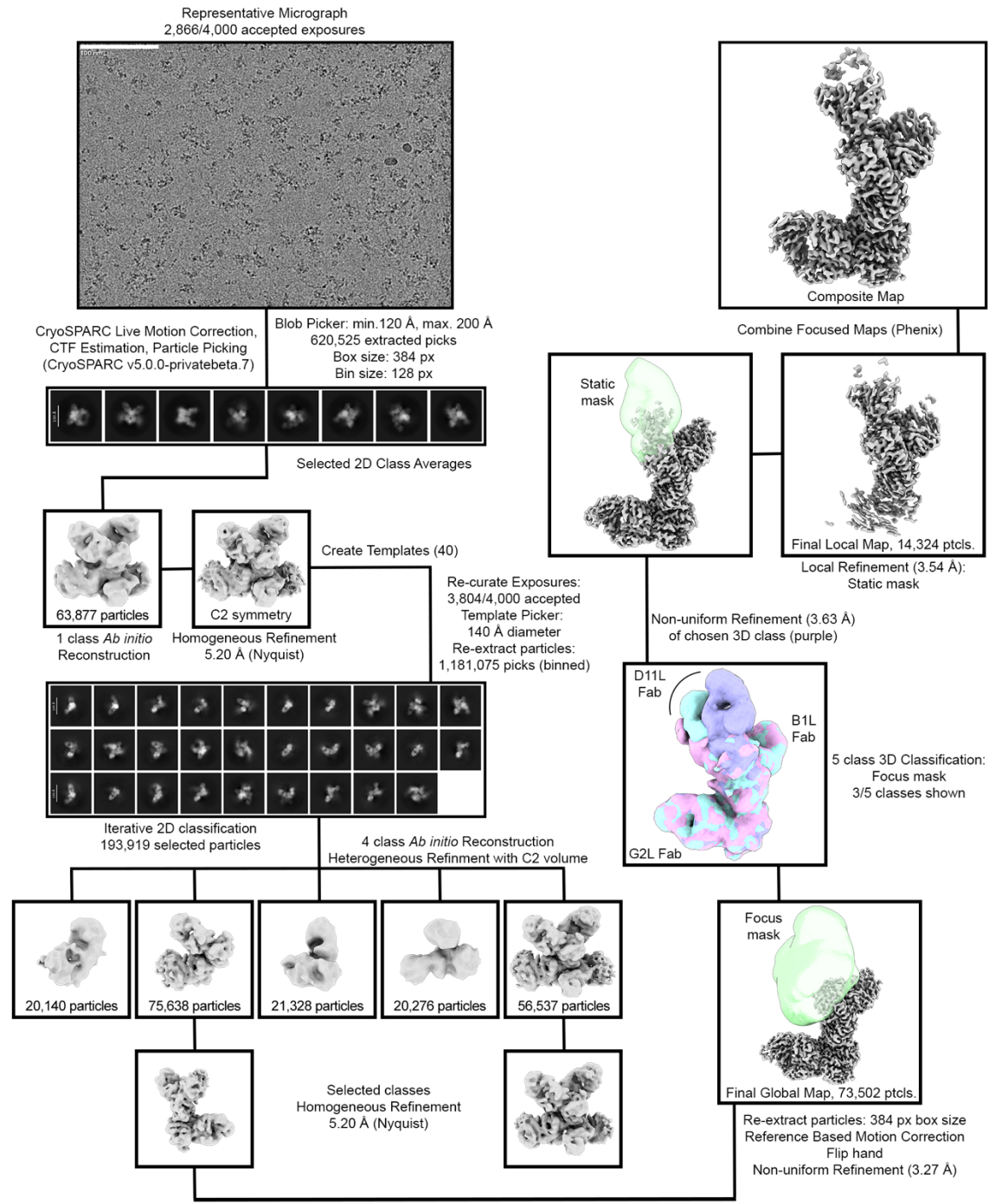

**Figure S4. Cryo-EM data processing workflow.**

Cryo-electron microscopy data processing workflow for HCMV gH/gK bound to Fabs D11L, B1L, and G2L. Connections between outputs indicate the selections made at each stage.

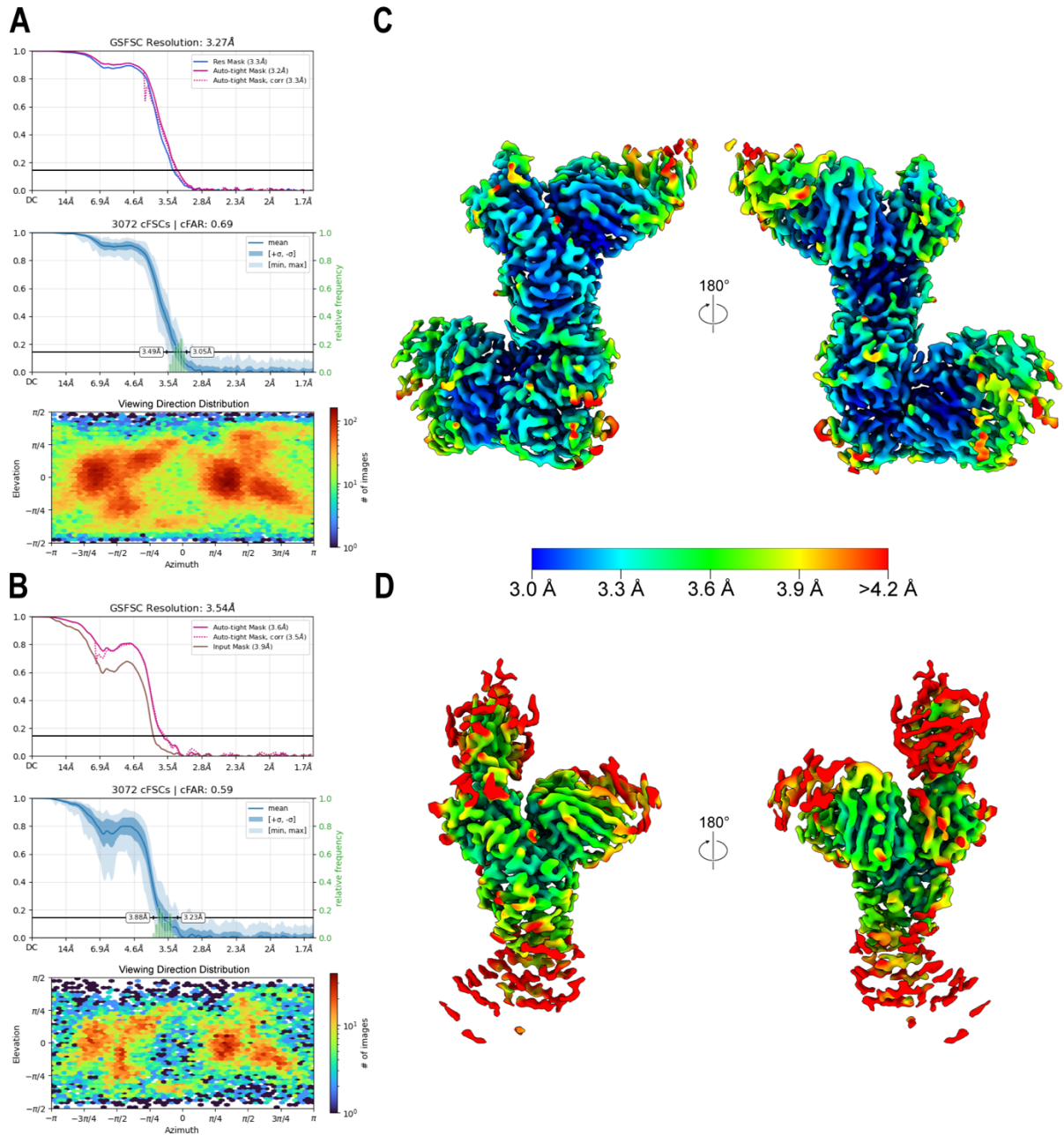

**Figure S5. Cryo-EM map validation.**

(A–B) Fourier shell correlation, conical Fourier shell correlation, and viewing direction distribution plots for the HCMV gH/gK + D11L + B1L + G2L global and local refinement maps, respectively. (C–D) Global and local refinement sharpened maps colored by local resolution, respectively. Two side views shown for each, rotated by 180 degrees.

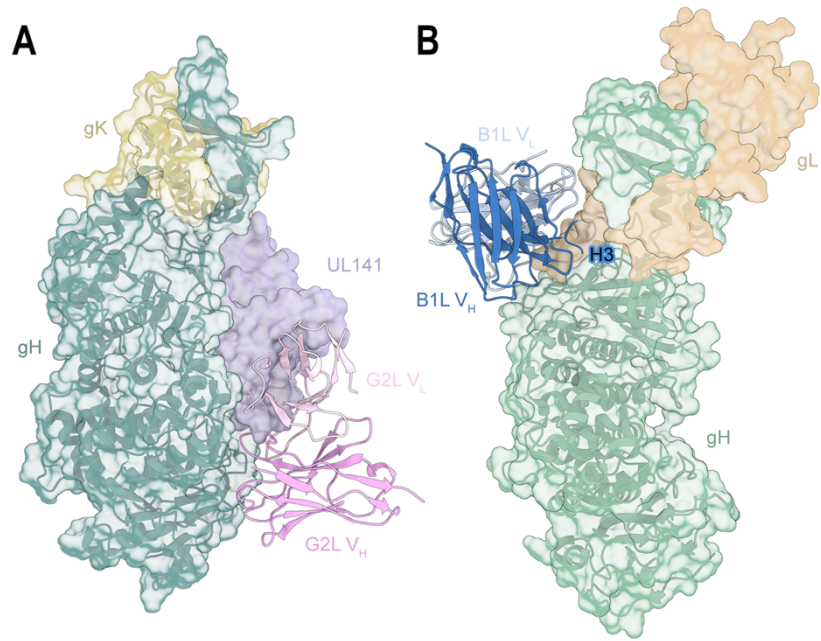

**Figure S6. HCMV gH/gK vs GATE and Trimer.**

(A) G2L docked onto HCMV GATE complex monomer (PDB: 9DIX, aligned by gH). GATE subunits (gH/gK/UL141) are shown as cartoons and transparent surfaces. Cartoon omitted for UL141 to show clash with G2L V<sub>L</sub>. (B) B1L docked onto HCMV Trimer (PDB: 8TCO, aligned by gH; gO hidden). Trimer subunits (gH/gL) are shown as cartoons and transparent surfaces. Cartoon omitted for gL to show clash with B1L CDRH3.

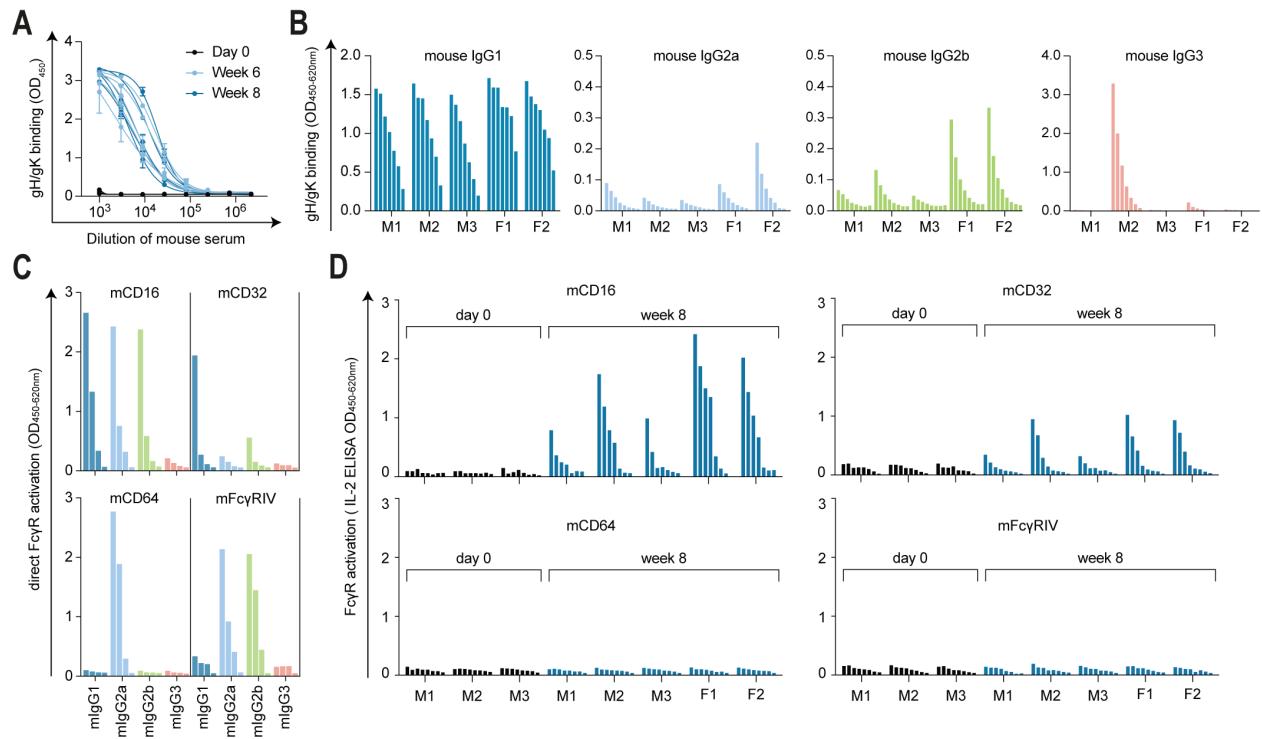

**Figure S7. Additional characterization of gH/gK-reactive antibody responses and FcγR activation in immunized mice.**

(A) ELISA-based detection of mouse IgG binding to the gH/gK complex in day 0, week 6, and week 8 sera from immunized mice. Optical density (OD) values are shown for each serum dilution, starting at 1:1,000 followed by 3-fold serial dilutions. Mean values and SDs from two separate experiments are shown. (B) Mouse IgG subclass-specific detection of gH/gK reactivity in week 8 sera from immunized mice, shown for each mouse (M1–M3 and F1–F2). Bound gH/gK-reactive antibodies were detected using mouse IgG subclass-specific antibodies. IgG subclasses are shown in different colors: mouse IgG1, dark blue; mouse IgG2a, light blue; mouse IgG2b, green; and mouse IgG3, orange. ELISA OD values are shown for each serum dilution, starting at 1:1,000 followed by 2-fold serial dilutions. (C) Direct FcγR activation by commercially available recombinant mouse IgG subclass control antibodies measured using the BW reporter system for each receptor: mCD16, mCD32, mCD64, and mFcγRIV. Mouse IgG subclass control antibodies were immobilized on plates at a starting concentration of 1 μg/ml followed by 2-fold serial dilutions. FcγR activation was measured by ELISA-based IL-2 detection in supernatants of the respective reporter cells, and ELISA OD values are shown. IgG subclasses are shown in the same color code as in Figure S7B. Each antibody was tested in two separate experiments; representative data from one experiment are shown. (D) Mouse FcγR activation by sera from immunized mice measured for each receptor (mCD16, mCD32, mCD64, and mFcγRIV) using the BW reporter system under gH/gK complex coating conditions. Activation was measured by ELISA-based IL-2 detection in supernatants of the respective reporter cells. ELISA OD values are shown for day 0 and week 8 serum samples, starting at a 1:1,000 dilution followed by 2-fold serial dilutions. Each sample was tested in two separate experiments; representative data from one experiment are shown.

| Sample | HCMV gH/gK + D11L, B1L, G2L Fabs |  |
| --- | --- | --- |
| Cryo-EM data collection |  |  |
| Microscope | FEI Titan Krios |  |
| Voltage (kV) | 300 |  |
| Detector | Gatan K3 |  |
| Energy filter | Gatan Biocontinuum |  |
| Slit width (eV) | 20 |  |
| Magnification (nominal) | 105,000 |  |
| Pixel size (Å/pix) | 0.8332 |  |
| Exposure rate (e<sup>-</sup>/pix/sec) | 17.6 |  |
| Exposure (e<sup>-</sup>/Å<sup>2</sup>) | 74.3 |  |
| Defocus range (μm) | 1.5–3.0 |  |
| Tilt angle (°) | 0 |  |
| Micrographs collected | 4,000 |  |
| Micrographs used | 3,804 |  |
| Particles extracted | 1,181,075 |  |
| Automation software | SerialEM |  |
| 3D reconstruction statistics |  |  |
| Refinement | Global | Local |
| Particles | 73,502 | 14,324 |
| Symmetry | C1 | C1 |
| Map sharpening B-factor | -101.5 | -64.6 |
| Resolution (Å) at FSC (half-maps) |  |  |
| Unmasked: 0.5 | 7.6 | 13.3 |
| Masked: 0.5 | 3.7 | 4.0 |
| Unmasked: 0.143 | 3.9 | 7.8 |
| Masked: 0.143 | 3.3 | 3.5 |
| EMDB ID | 79522 | 79524 |
| Model refinement and validation statistics |  |  |
| Composition |  |  |
| Amino Acids | 1416 |  |
| Ligands (type) | 2 (NAG) |  |
| Bonds (RMSD) |  |  |
| Length (Å) (# > 4σ) | 0.003 (0) |  |
| Angles (°) (# > 4σ) | 0.57 (0) |  |
| Ramachandran plot |  |  |
| Outliers (%) | 0.0 |  |
| Allowed (%) | 4.0 |  |
| Favored (%) | 96.0 |  |
| Rotamer outliers (%) | 0.0 |  |
| C-β outliers (%) | 0.0 |  |
| CaBLAM outliers (%) | 0.96 |  |
| ADP (B-factors) |  |  |
| Amino Acids (mean) | 61.0 |  |
| Ligands (mean) | 73.7 |  |
| CC (mask) | 0.86 |  |
| MolProbity score | 1.39 |  |
| Clash score | 3.23 |  |
| EMRinger score | 3.47 |  |
| PDB ID | 39JB |  |
| EMDB ID (composite map) | 79526 |  |

**Table S1. Cryo-electron microscopy data collection and refinement statistics.**

Statistics for data collection, processing, model building, and refinement are shown.
